# The fate of recessive deleterious or overdominant mutations near mating-type loci under partial selfing

**DOI:** 10.1101/2022.10.07.511119

**Authors:** Emilie Tezenas, Tatiana Giraud, Amandine Véber, Sylvain Billiard

**Author notes:** Corresponding author, Unité Evo-Eco-Paleo, Bât. SN2, Cité Scientifique, 59655 Villeneuve d’Ascq, France. Co-senior author (Tatiana Giraud, Amandine Véber, Sylvain Billiard).

## Abstract

Large regions of suppressed recombination having extended over time occur in many organisms around genes involved in mating compatibility (sex-determining or mating-type genes). The sheltering of deleterious alleles has been proposed to be involved in such expansions. However, the dynamics of deleterious mutations partially linked to genes involved in mating compatibility are not well understood, especially in finite populations. In particular, under what conditions deleterious mutations are likely to be maintained for long enough near mating-compatibility genes remains to be evaluated, especially under selfing, which generally increases the purging rate of deleterious mutations. Using a branching process approximation, we studied the fate of a new deleterious or overdominant mutation in a diploid population, considering a locus carrying two permanently heterozygous mating-type alleles, and a partially linked locus at which the mutation appears. We obtained analytical and numerical results on the probability and purging time of the new mutation. We investigated the impact of recombination between the two loci and of the mating system (outcrossing, intra and inter-tetrad selfing) on the maintenance of the mutation. We found that the presence of a fungal-like mating-type locus (*i.e*. not preventing diploid selfing) always sheltered the mutation under selfing, *i.e*. it decreased the purging probability and increased the purging time of the mutations. The sheltering effect was higher in case of automixis (intra-tetrad selfing). This may contribute to explain why evolutionary strata of recombination suppression near the mating-type locus are found mostly in automictic (pseudo-homothallic) fungi. We also showed that rare events of deleterious mutation maintenance during strikingly long evolutionary times could occur, suggesting that deleterious mutations can indeed accumulate near the mating-type locus over evolutionary time scales. In conclusion, our results show that, although selfing purges deleterious mutations, these mutations can be maintained for very long times near a mating-type locus, which may contribute to promote the evolution of recombination suppression in sex-related chromosomes.

## 1 Introduction

The evolution of sex chromosomes, and more generally of genomic regions lacking recombination, is widely studied in evolutionary biology as it raises multiple, unresolved questions (Ironside, 2010, Yan et al., 2020, Hartmann, Ament-Velásquez, et al., 2021, Kratochvíl and Stöck, 2021, Jay et al., 2022). A striking feature of many sex and mating-type chromosomes is the absence of recombination in large regions around the sex-determining genes. Recombination suppression indeed evolved in various groups of plants and animals in several steps beyond the sex-determining genes, generating evolutionary strata of differentiation between sex chromosomes (Nicolas et al., 2004, Bergero and Charlesworth, 2009, Hartmann, Duhamel, et al., 2021, Kratochvíl and Stöck, 2021). The reasons for the gradual expansion of recombination cessation beyond sex-determining genes remain debated (Ironside, 2010, Wright et al., 2016, Ponnikas et al., 2018, Hartmann, Duhamel, et al., 2021). Recombination suppression has extended progressively with time not only on many sex chromosomes but also on mating-type chromosomes in fungi (Hartmann, Duhamel, et al., 2021) and other supergenes (Yan et al., 2020, Jay et al., 2021).

The main hypothesis to explain such stepwise extension of recombination cessation on sex chromosomes has long been sexual antagonism (Charlesworth et al., 2005, Bergero and Charlesworth, 2009). Theoretical studies have indeed shown that the suppression of recombination may evolve to link alleles that are beneficial in only one sex to the sex-determining genes (Rice, 1987, Charlesworth et al., 2005, Ruzicka et al., 2020). However, this hypothesis has received little evidence from empirical studies despite decades of research (Ironside, 2010, Dagilis et al., 2022). Moreover, the sexual antagonism hypothesis cannot explain the evolutionary strata found on fungal mating-type chromosomes. Indeed, in many fungi, two gametes can form a new individual only if they carry different mating types, but there is no sexual antagonism or other form of antagonistic selection between cells of opposite mating types; the cells of different mating types do not show contrasted phenotypes or footprints of diversifying selection (Bazzicalupo et al., 2019). Yet, evolutionary strata have been documented on the mating-type chromosomes of multiple fungi, with recombination suppression extending stepwise beyond mating-type determining genes (Fraser et al., 2004, Menkis et al., 2008, Branco et al., 2017, Branco et al., 2018, Hartmann, Duhamel, et al., 2021, Hartmann, Ament-Velásquez, et al., 2021, Vittorelli et al., 2022). Evolutionary strata have also been reported around other supergenes, *i.e*., large genomic regions encompassing multiple genes linked by recombination suppression, such as in ants and butterflies (Yan et al., 2020, Jay et al., 2021). Several hypotheses alternative to sexual antagonism have been proposed and explored to explain the stepwise extension of recombination suppression on sex-related chromosomes (Ironside, 2010, Hartmann, Duhamel, et al., 2021). Theoretical models suggested that recombination suppression could be induced by a divergence increase in regions in linkage disequilibrium with a sex-determining locus (Jeffries et al., 2021) or that inversions could be stabilized by dosage compensation on asymmetric XY-like sex chromosomes (Lenormand and Roze, 2022).

A promising, widely applicable hypothesis is the sheltering of deleterious alleles by inversions carrying a lower load than average in the population (Charlesworth and Wall, 1999, Antonovics and Abrams, 2004, Hartmann, Duhamel, et al., 2021, Jay et al., 2022). Inversions (or any suppressor of recombination in *cis*) can indeed behave as overdominant: inversions with fewer recessive deleterious mutations than average are initially beneficial and increase in frequency, but can then occur in a homozygous state where they express their load, unless they are linked to a permanently heterozygous allele. In this case, they remain advantageous, and can reach fixation in the sex-related chromosome on which they appeared (Jay et al., 2022). The suppression of recombination is thereby selected for, and recessive deleterious mutations are permanently sheltered. The process can occur repeatedly, leading to evolutionary strata. Importantly, this is one of the few hypotheses able to explain the existence of evolutionary strata on fungal mating-type chromosomes and it can apply to any supergene with a permanently heterozygous allele (Llaurens et al., 2017, Jay et al., 2022).

A key point for the recombination suppressor to invade is that it must appear in populations where recessive deleterious mutations segregate near the mating-compatibility genes (Olito et al., 2022, Jay et al., 2022). We therefore need to understand whether such mutations can persist in the vicinity of permanently heterozygous alleles (such as those occurring at mating-type loci) and under what conditions. In particular, it is usually considered that selfing purges deleterious mutations (Glémin, 2007, Abu Awad and Billiard, 2017), while most evolutionary strata on fungal mating-type chromosomes have been reported in selfing (automictic) fungi (Branco et al., 2017, Branco et al., 2018, Hartmann, Ament-Velásquez, et al., 2021, Vittorelli et al., 2022). Indeed, because mating types are determined at the haploid stage in fungi, mating types do not prevent selfing when considering diploid individuals (Billiard et al., 2012). Some particular forms of selfing associated with a permanently heterozygous mating-type locus such as intra-tetrad mating (*i.e*. automixis, mating among gametes from the same meiosis) can however favor the maintenance of heterozygosity (Hood and Antonovics, 2000). Indeed, mating can only occur between haploid cells carrying different mating-type alleles, which maintains heterozygosity at the mating-type locus, and to some extent at flanking regions, thereby possibly sheltering deleterious alleles. We therefore need to study whether deleterious or overdominant mutations can be maintained near mating-type compatibility loci, even under selfing, to assess whether the mechanism of sheltering deleterious mutations can drive extensions of recombination suppression.

The dynamics of deleterious mutation frequencies in genomes have been extensively studied independently of the presence of a permanently heterozygous locus. Deterministic models and diffusion approximations have been used to study the dynamics of deleterious mutations in a one locus-two allele setup (Kimura, 1980, Ewens, 2004, Rice, 2004), with the addition of sexual reproduction and in particular selfing (Ohta and Cockerham, 1974, Caballero and Hill, 1992, Abu Awad and Roze, 2018). Extensions of these models exist to cover the two locus-two allele case (Karlin, 1975) and multilocus systems (reviewed in Bürger, 2020), or to take stochastic fluctuations into account (Coron et al., 2013, Coron, 2014). However, the dynamics of deleterious mutations in genomic regions near a permanently heterozygous allele have been little studied. A deterministic model showed that a lethal allele can be sheltered in an outcrossing population only when it is completely linked to a self-incompatibility locus (Leach et al., 1986). Another deterministic model introduced selfing and showed with simulations that a lethal allele can be sheltered when it is completely linked to a mating-type allele, favored in a heterozygote state, and if there is intra-tetrad selfing (Antonovics et al., 1998). Assuming a variable recombination rate between the two loci, Antonovics and Abrams, 2004 showed that an overdominant allele lethal in a homozygous state could be maintained if recombination was twice as low as the selection for heterozygotes and mating occurred via intra-tetrad selfing. Stochastic simulations additionnally showed that a recessive deleterious allele could be maintained completely linked to a self-incompatibility allele, especially when it is highly recessive, and when the number of self-incompatibility alleles in the population is large (Llaurens et al., 2009), and that codominant weakly deleterious alleles could be maintained near loci under balancing selection in the major histocompatibility complex (MHC) in humans (Lenz et al., 2016).

Here, building on the work of Antonovics and Abrams, 2004, we use a similar though simplified two locus-two allele framework, taking into account the non-negligible reproductive stochasticity during the early stage of the dynamics of the mutant subpopulation, until it becomes extinct or reaches some appreciable fraction of the total population. More precisely, we consider a permanently heterozygous mating-type locus and a genetic load locus, and we assume that the recombination rate between the two loci is a fixed parameter. Individuals can reproduce via outcrossing, or via either one of two types of selfing, intra-tetrad mating or inter-tetrad mating. The two types of selfing depend on whether a given gamete mates with another gamete produced during the same meiosis event (within a tetrad) or with a gamete from a different meiosis (from another tetrad, App. **??**). The distinction is important because intra-tetrad mating maintains more heterozygosity in some genomic regions than inter-tetrad mating (Hood and Antonovics, 2000). Starting with a continuous-time Moran process, we derive the rates at which individuals of each genotype are produced. Then, as a new mutation is carried by very few individuals at the beginning of its evolution, a branching process naturally arises. Indeed, in this initial phase two individuals carrying the mutant allele have an extremely low probability to mate with each other. Mutant-carrier individuals can thus be assumed to reproduce independently of each other, leading to an approximation of the dynamics of the subpopulation of mutant carriers by a branching process.

The use of branching processes has shown its utility to account for the dynamics of a newly arised mutant allele in a population. Many estimates of the fixation or purging time of mutants in stochastic models (Champagnat and Méléard, 2011, Collet et al., 2013) relied on the use of branching processes to approximate the dynamics of a newly appeared mutant allele and of a nearly-fixed one. A branching-process approximation was used to study a two locus-two allele model, with individual fitness depending on the allelic state at both loci (Ewens, 1967, Ewens, 1968). For the diploid case, the framework of a seven-type branching process that can be used to study the fate of a deleterious mutation has been described, without deriving any analytical result (Pollard, 1966, Pollard, 1968). A similar branching process approximation was used to study the fate of a beneficial mutation with selfing (Pollak, 1987, Pollak and Sabran, 1992). Here, we use a similar framework but consider deleterious mutations and a permanently heterozygous locus. Modeling multiple loci suggests the use of multitype branching processes, which have been widely studied (Harris, 1964, Kesten and Stigum, 1966, Mode, 1971, Athreya and Ney, 1972, Sewastjanow, 1975, Pénisson, 2010). However, the multiplicity of types renders the derivation of analytical results on probabilities of extinction and on extinction times difficult (Heinzmann, 2009). We therefore use an analytical approach to study the probability that a new mutation is purged from the population, and a numerical approach to study the purging time (when purging occurs) to assess how long a deleterious or overdominant mutation remains in a population. We study in particular the impact of the mating system and of the level of linkage to a permanently heterozygous locus on the long-term maintenance of deleterious mutations near a fungal-like mating-type locus (*i.e*. not preventing diploid selfing).

## 2 Methods and Models

All parameters which will be needed below are listed in App. **??**.

### 2.1 Population and stochastic dynamics

We consider diploid (or dikaryotic) individuals, represented by their mating-type chromosomes, that harbor two biallelic loci: one mating-type locus, with alleles *A* and *a*, and one load locus, with a wild allele *B* and a mutant allele *b*. We model a fungal-like mating-type locus, so that mating is only possible between haploid cells carrying different alleles at the mating-type locus (this does not prevent diploid selfing as each diploid individual is heterozygous at the mating-type locus). Consequently, only four genotypes are admissible, denoted by *G*_1_, …, *G*_4_ in Figure **??**. We follow the evolution of (*g*(*t*)_*t≥*0_ = (*g*_1_(*t*), …, *g*_4_(*t*))_*t≥*0_, where *g*_*i*_(*t*) is the number of individuals of genotype *G*_*i*_ in the population at time *t*. We suppose that the reproduction dynamics is given by a biparental Moran model with selection. In this continous-time model, a single individual is replaced successively and the total population size, denoted by *N*, remains constant. A change in the population state *g* occurs in three steps.

The first step is the production of an offspring. After a random time following an exponential law of parameter *N*, an individual is chosen uniformly at random to reproduce. This means in particular that all individuals have the same probability to reproduce. Mathematically speaking, this formulation is equivalent to saying that each individual reproduces at rate 1. The chosen diploid individual produces haploid gametes, via meiosis, during which recombination takes place between the two loci with probability *r* (see Figure **??** (a)). The product of a meiosis is a tetrad that contains four haploid gametes (Figure **??** (b)). Mating can then occur through three modalities, illustrated in App. **??** (recall that two gametes can fuse only if they carry different mating-type alleles): (i) Intra-tetrad selfing, with probability *fp*_*in*_: the two gametes are picked from the same tetrad, only one parent is involved; (ii) Inter-tetrad selfing, with probability *f* (1 − *p*_*in*_): the two gametes are picked from two different tetrads produced by the same individual, only one parent is involved; (iii) Outcrossing, with probability 1 − *f* : the two gametes are picked from tetrads produced by two different parents. In this case, the second parent is chosen uniformly at random in the remaining population, and produces haploid gametes via meiosis with the same recombination rate *r*. An offspring is produced following the chosen mating system, its genotype thus depending on the genotypes of the parents involved and on the occurrence of a recombination event in the tetrads.

The second step is the offspring survival. We assume that the fitness of a genotype *G*_*i*_ is the probability that an offspring with that genotype survives, and we denote it by *S*_*i*_. We consider two selection scenarii (Figure **??**, left): (i) The partial dominance case, where the mutant allele *b* is always deleterious and recessive. Homozygotes *bb* and heterozygotes *Bb* at the load locus have fitness values (*i.e*. a probability of survival) of 1 − *s* and 1 − *hs*, respectively. Homozygotes *BB* have fitness 1; (ii) The overdominance case, where heterozygotes *Bb* are favored over *BB* and *bb* individuals. In this case, the fitness of *Bb, bb* and *BB* juveniles are respectively 1, 1 − *s*_3_ and 1 − *s*_4_, with *s*_3_ > *s*_4_ so that the fitness of *bb* individuals is lower than the fitness of wild-type individuals *BB*. The mating-type locus is considered neutral regarding survival.

The third step occurs if the offspring survives, in which case an individual chosen uniformly at random in the extant population is chosen to die and to be replaced by the offspring. If the offspring does not survive, the population state (*g*_1_, *g*_2_, *g*_3_, *g*_4_) does not change.

A jump in the stochastic process is thus an increase by one of the number of genotype *G*_*i*_ individuals in the population, when an offspring of genotype *G*_*i*_ is produced and survives, and a concomitant decrease by one of the number of genotype *G*_*j*_ individuals in the population, when an adult of genotype *G*_*j*_ dies. If *i* = *j, i.e*. if the surviving offspring and the individual chosen to die have the same genotype, the composition of the population does not change. We denote the jump rate from *g* to *g* + *e*_*i*_ − *e*_*j*_ by *Q*_*i,j*_(*g*), where *e*_*i*_ is the vector with a 1 in position *i* and zeros everywhere else. *Q*_*i,j*_(*g*) is equal to the product of the rate at which an offspring of genotype *G*_*i*_ is produced (first step), which we denote by *T*_*g*_(+*G*_*i*_), of the probability that it survives (*S*_*i*_, second step), and of the probability that the adult chosen to die is of genotype *G*_*j*_ (third step). Thus, we have

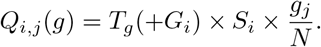

The total rates at which individuals of different genotypes are produced are given in App. **??**. For example, the rate at which an offspring of genotype *G*_1_ is produced when the current state of the population is *g* = (*g*_1_, *g*_2_, *g*_3_, *g*_4_) is given by

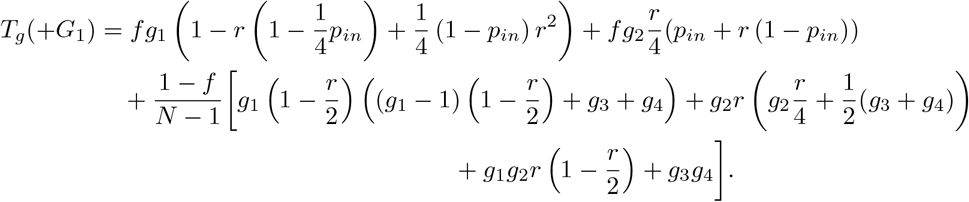

The first two terms on the right-hand side, with a factor *f*, correspond to reproduction events by selfing. The third term, with a factor 1 − *f*, corresponds to reproduction events by outcrossing. Each subterm then encompasses the rate at which each genotype is involved in the reproduction event, and the probability that the offspring produced is of genotype *G*_1_, taking into account possible recombinations. For example, the subterm (1 − *f*)/(*N* − 1) × *g*_1_(*g*_1_ − 1)(1 − *r*/2)^2^ is the product of the total rate *g*_1_ × 1 at which an individual of genotype 1 reproduces, of the probability 1 − *f* that reproduction happens by outcrossing, of the probability (*g*_1_ − 1)/(*N* − 1) that the second parent is chosen among the other individuals of genotype *G*_1_, and of the probability (1 − *r*/2)^2^ that their offspring has genotype *G*_1_.

**Figure 1:**
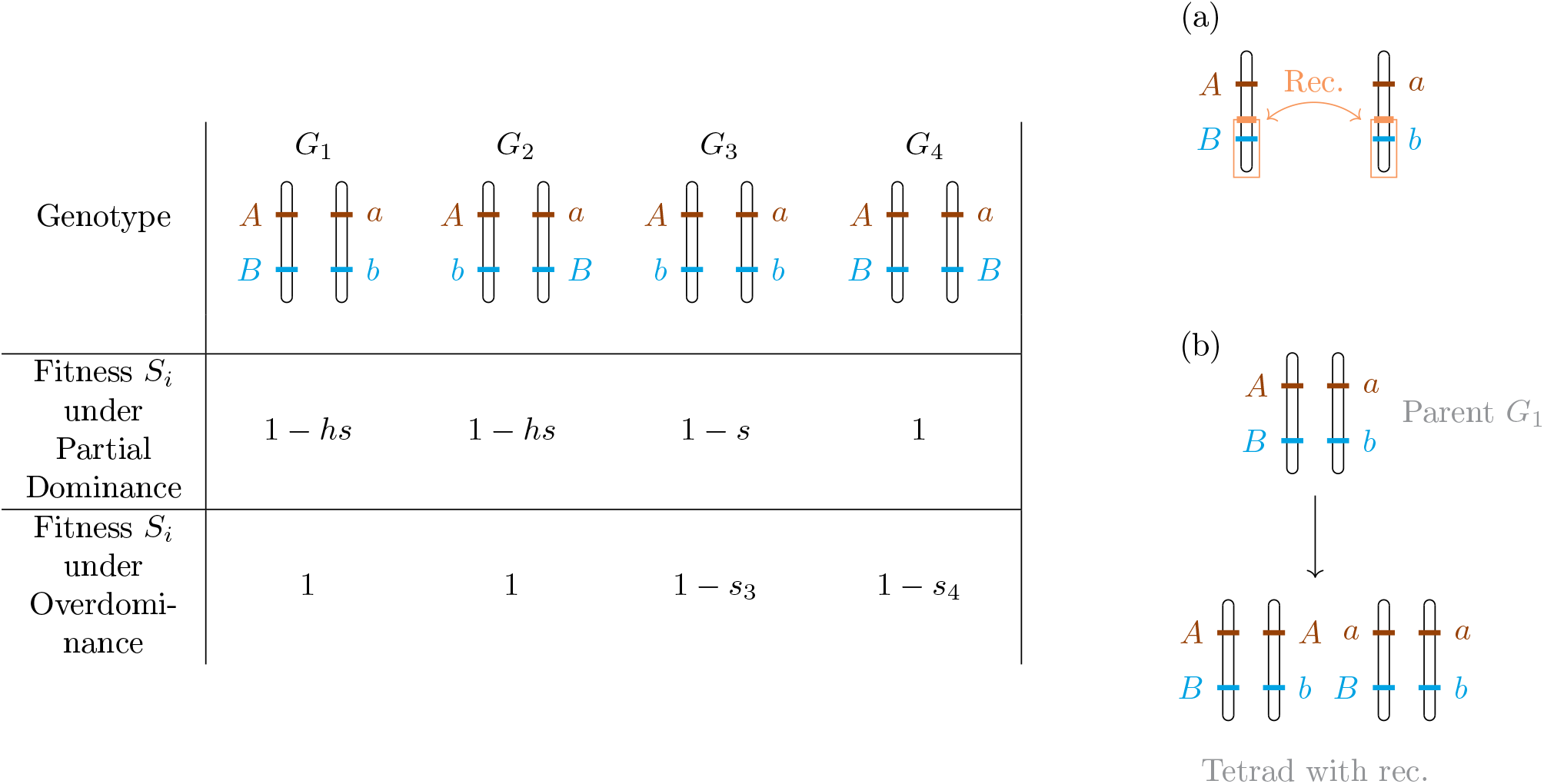
Schematic drawings of the genotypes considered and their parameters. (Left) Description of the possible genotypes in the population and their fitness *S*_*i*_ for the two selection scenarii considered (partial dominance and overdominance). (Right) (a) Position of a putative event of recombination between the mating-type locus and the load locus. (b) Example of a tetrad that can be obtained after a meiosis of an individual of genotype *G*_1_, with recombination. Four gametes are produced, two of each mating type. In the second and third gamete from the left, combinations of alleles that did not exist in the parent are observed (*A* with *b* and *a* with *B*).

### 2.2 Branching-Process approximation

Let us now consider that the population size *N* is very large. When a mutation appears at the load locus, it is carried by a single individual. Hence, during the initial phase of the dynamics of the mutation *b*, the number of individuals who carry the mutation remains small compared to the number of wild-type individuals. The number of wild-type individuals is of the same order of magnitude as the total population size *N*, and the number of mutation-carrier individuals is negligible. More precisely, we assume that, when *N* is large,

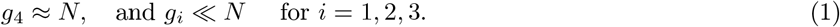

Under this assumption, the jump rates *Q*_*i,j*_(*g*) of the process can be approximated by neglecting the terms of the form 1/(*N* − 1) × *g*_*i*_ × *g*_*j*_, with *i, j* ∈ {1, 2, 3}, as they are of order 1*/N*. This means that mating by outcrossing between individuals carrying the mutation *b* can be neglected. As a consequence, the birth rates of the different genotypes are linear in *g*_*i*_, and a reproduction law for each genotype that is independent of the number of individuals of all other mutant-carrier genotypes can be derived. The Moran process can then be approximated by a branching process that follows the change in genotype counts for the mutation-carrier genotypes only.

We denote this branching process by (*Z*_*t*_)_*t≥*0_, where for each *t* ≥ 0, we have *Z*_*t*_ = (*Z*_*t*,1_, *Z*_*t*,2_, *Z*_*t*,3_), with *Z*_*t,i*_ the number of individuals of genotype *G*_*i*_ in the population at time *t*. To each genotype is associated a reproduction law, that is, a probability distribution on ℕ^3^ (vectors with three integer-valued coordinates) that gives the probability for an individual of that genotype to produce a given number of descendants of each genotype when it reproduces. Note that the rationale behind the branching process is different from the one for the Moran process. Indeed, each *replacement event* in the Moran model that involves an individual carrying the mutant allele *b* will be seen in the branching process as a *reproduction event*, in which the *offspring* is the mutant individual that is possibly produced during the first step of the Moran jump, and the *parent* is another mutant individual that is either one of the two actual parents in the replacement event, or the individual chosen to be replaced by the offspring in the Moran replacement event. A *reproduction event* of the branching process consists in the replacement of the parent by its descendants, which will be made of the mutant *offspring* when there is one, and of the mutant *parent* when it remains in the population. More precisely, we will encode three situations as follows: (i) when the replacement event in the Moran model corresponds to the reproduction of an individual of genotype *G*_*i*_, *i* ∈ {1, 2, 3} (via selfing or outcrossing with an individual of genotype *G*_4_), that this reproduction event generates a mutant offspring of genotype *G*_*j*_, *j* ∈ {1, 2, 3}, and the mutant parent is not chosen to die, we will see the *reproduction event* of the branching process as being an individual of genotype *G*_*i*_ having descendance vector *e*_*i*_ + *e*_*j*_; (ii) When the Moran replacement event leads to the reproduction of an individual of genotype *G*_*i*_, *i* ∈ {1, 2, 3} (via selfing or outcrossing with an individual of genotype *G*_4_), that this reproduction event generates an offspring of genotype *G*_4_, and the mutant parent is not chosen to die, we will see the *reproduction event* as being an individual of genotype *G*_*i*_ having descendance vector *e*_*i*_ (as non-mutant individuals are not accounted for in the branching process approximation).

Note that this reproduction event will imply no change in the population state, but for the sake of completeness we indicate here all Moran replacement events that have non-vanishing rates as *N* tends to infinity; (iii) When the Moran replacement event only involves non-mutant parents and an individual of genotype *G*_*i*_, *i* ∈ {1, 2, 3}, is chosen to die, we will see the *reproduction event* as being an individual of genotype *G*_*i*_ having descendance vector 0 (corresponding to the *parent* being removed from the branching process and no mutant offspring being produced). Other possible Moran replacement events occur at rates that vanish as *N* tends to infinity, and therefore do not contribute to the *reproduction events* of the branching process. The rates at which *reproduction events* described above occur are directly derived from the rates *Q*_*i,j*_(*g*) of the Moran model, under the approximation stated in Eq.(**??**). They are summarized in the matrices *A, T*, and *D* defined as follows:

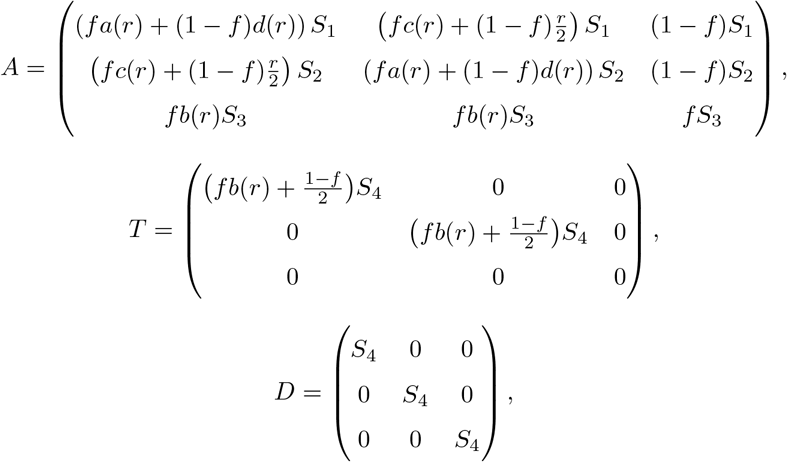

with

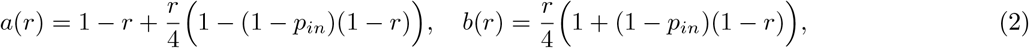

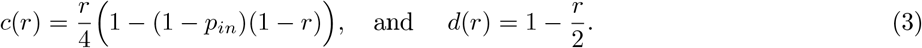

The entries *A*_*ij*_ of matrix *A, T*_*ij*_ of matrix *T* and *D*_*jj*_ of matrix *D* give the rates at which each individual of genotype *j* reproduces and gives rise to a descendance vector respectively equal to *e*_*i*_ + *e*_*j*_ (situation (i)), *e*_*i*_ (situation (ii)), and 0 (situation (iii)). An example of derivation of the matrix coefficients is given in App. **??**.

### 2.3 Probability of purge and purging time

Under the assumption that the mutation is initially rare (after a mutation or migration event for example), we can use the branching process approximation described in Section **??** to derive the probability and purging time of the mutation from the population. In particular, our goal is to analyze the effect of the presence of a mating-type locus near the load locus on the purge of the deleterious mutant *b, i.e*. on the extinction of the mutant-carrier population described by the branching process.

#### Extinction Probability

The probability of extinction of the branching process can be determined by looking at the eigenvalues of the matrix *C* such that 𝔼[*Z*_*t*_|*Z*_0_ = *z*_0_] = *z*_0_*e*^*Ct*^ for *t* ≥ 0, where *z*_0_ ∈ ℕ^3^ is the initial state of the branching process (*Z*_*t*_)_*t≥*0_ (Sewastjanow, 1975 in German, and Pénisson, 2010 for a statement of these results in English). Under the assumption of irreducibility of the matrix *C*, results relying on the theory of Perron-Froebenius (see for example Athreya and Ney, 1972) state that the process almost surely dies out (*i.e*. the mutation is purged with probability 1) if and only if *ρ*, the maximum eigenvalue of *C*, satisfies *ρ* ≤ 0. When *C* is not irreducible, which occurs for example if *f* = 0 or *f* = 1, the result still holds but requires the use of the theory of final classes (Sewastjanow, 1975, cited in Pénisson, 2010). Details are given in App. **??**.

We follow a method described in Bacaër, 2018, to compute the matrix *C* mentioned above and obtain

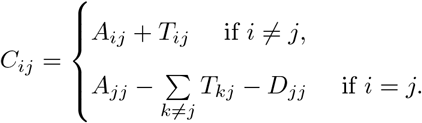

This gives

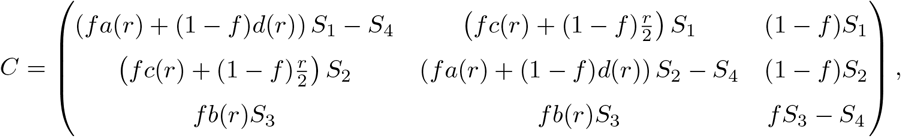

where the functions *a, b, c, d* were defined in Eqs. **??** and **??** (see details in App. **??**).

We derived the dominant eigenvalue using Mathematica (Wolfram Research, 2015) and study its sign analytically when possible, or numerically otherwise.

#### Comparison with previous results

Our results can be compared to the work of Ewens, 1967, who used a similar framework to study a random-mating population with two biallelic loci under selection, one of which carried a new allele. Assuming that the frequency of the gametes that carried a new allele was negligible compared to the frequencies of wild-type gametes, he used a branching process approximation to study the probability that the new allele was purged from the population. He considered a recombination rate *R* between the two loci, and fitnesses *w*_*ij*_ for each genotype (where *i* and *j* take the value 1 or 3 when loci are homozygous, and the value 2 when heterozygous). Setting *w*_*i*1_ = *w*_*i*3_ = 0 for *i* = 1, 2, 3 allows to force heterozygosity at the locus that does not carry the new allele in his model, and to compare his findings with our results on the fate of a new allele appearing near a permanently heterozygous locus. The dominant eigenvalue of the matrix driving the dynamics of the new allele in Ewens, 1967, is

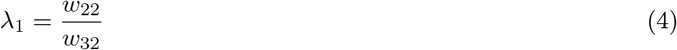

with *w*_22_ being the fitness of individuals heterozygous for the new allele, and *w*_32_ the fitness of homozygous wild-type individuals. As Ewens, 1967, considered a discrete-time branching process, this dominant eigenvalue must be compared to one to deduce information on the new allele survival probability.

#### Sheltering effect of the mating-type locus

We investigate now to the potential effect of the presence of a mating-type locus on the maintenance of a mutant allele in a population: as mating-type alleles are always heterozygous, any mutation appearing completely linked to one mating-type allele is maintained in a heterozygous state as well. The load of the mutant allele is then less expressed when the mutation is recessive, and the mutation is said to be “sheltered”.

This potential *sheltering effect* can be explored by looking at the variation of the dominant eigenvalue *ρ* when the recombination rate *r* is close to 0.5. Indeed, the quantity |*ρ*| can be seen as the rate of decay of the deleterious mutant subpopulation (see the results on the probability of survival of a multitype branching process, Th. 3.1 of Heinzmann, 2009), and its value gives a rough approximation of the inverse of the mean time to extinction of this subpopulation, *i.e*. of the mean purging time of the mutant allele *b*. Moreover, setting the recombination rate to *r* = 0.5 in our model allows us to consider a load locus completely unlinked to the mating-type locus, while decreasing the value of *r* introduces some loose linkage between the two loci. We thus look at the derivative 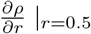 to obtain the variation of the dominant eigenvalue of *C* when departing from this unlinked state.

The sign of the derivative gives information on the existence of a sheltering effect due to the mating-type locus: 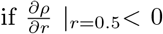, then when *r* decreases from 0.5 to lower values, *i.e*. when linkage between the two loci appears, the (negative) value of *ρ* increases, which means that the purging of the mutation becomes slower. In this case, the mating-type locus has a sheltering effect. Otherwise, if 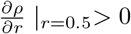, the presence of a mating-type locus accelerates the purging of a deleterious allele.

The absolute value of the derivative also gives information on the strength of the sheltering effect of the mating-type locus. The closer to 0 the derivative is, the smaller the impact of the mating-type locus. We compute the derivative and study its sign analytically. We then study the values of the derivative numerically in order to identify the impact of each parameter on the sheltering effect of the mating-type locus.

We also look at the strength of the sheltering effect on mutations close to the mating-type locus, by studying the eigenvalue variation around *r* = 0. Setting the recombination rate to *r* = 0 models a situation where the load locus is completely linked to the mating-type locus. Hence, the mutation is completely linked to one mating-type allele, and maintained in a heterozygous state. Looking at the derivative 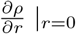 allows us to quantify the impact of departing from this situation by loosening the linkage between the two loci. We study the difference between the derivative at *r* = 0.5 and the derivative at *r* = 0 to compare the effect of adding a small amount of linkage between completely unlinked loci (*r* = 0.5) and the effect of adding a small amount of recombination between completely linked loci (*r* = 0).

#### Extinction time

The mean time to extinction in a multitype branching process is finite for a subcritical process (that is, when the principal eigenvalue *ρ* of *C* is less than 0), and infinite for a critical process (*i.e*. when *ρ* = 0, see Pötscher, 1985, for the proof of existence and finiteness of extinction time moments). Previous work, in particular Theorem 4.2 in Heinzmann, 2009, showed that a Gumbel law gives a good approximation of the law of the extinction time, provided that the initial number of individuals in the branching process and the absolute value of the dominant eigenvalue are both large. In our case, however, the mutation appears in a single individual, and the dominant eigenvalue is close to zero, which prevents the use of the Gumbel law approximation. Therefore, we performed computer simulations to study the empirical distribution of the time to extinction of the process, *i.e*. the purging time of the *b* mutant allele.

The branching process was simulated with a Gillespie algorithm to obtain an empirical distribution for the time to extinction. More precisely, the Gillespie algorithm produces realizations of the stochastic process by iteratively updating the number of individuals of each genotype within the multitype branching process (Gillespie, 1976). To circumvent the problem of exponential increase of the population size in the supercritical case, the parameters were chosen so that the branching process was subcritical. The probability of extinction was thus equal to 1 and the mean time to extinction was finite. For each scenario, we looked at different values of the recombination rate *r*, in order to study the impact of linkage between the load locus and the mating-type locus on the purging time of the mutant allele. We also chose different values for the selfing rate *f* in order to assess the impact of the mating system on the purging time of the mutant allele. For each set of parameters, 100,000 independent simulation runs were performed with the same initial condition (a single individual heterozygous at the load locus was introduced).

#### Probability of a new mutation apparition before the first one is purged

As a first step towards the study of the accumulation of deleterious mutations near a mating-type locus, we studied the probability that the deleterious mutation can be maintained long enough in the population so that a second mutation can appear before the first one is purged. We considered that a second mutation could appear during a reproduction event occurring in the population of mutation carriers (described by the branching process), on a region of a given length *d* = 10^6^ base pairs, at a rate of *µ* = 10^−8^ mutations per base pairs per reproduction event. The mean number of reproduction events needed for a new mutation to appear in a region of length *d*, 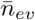, is the inverse of the mutation rate *µ* multiplied by the length *d*:

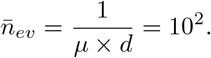

We then estimated the probability that a new mutation appears in such a genomic region before the first one is purged by counting the number of independent simulations in which the number of reproduction events exceeded 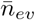 before the branching process went extinct (*i.e*. before the purging of the first mutation), over 100,000 simulation runs. Note that we did not take into account the genotype of the individual on which the second mutation appears, and therefore we did not distinguish whether the second mutation appears on a chromosome that carries the first one or not. Our estimate thus does not exactly equals the probability to have two mutations on the same chromosome, but this gives an order of magnitude of the probability of deleterious mutation accumulation and of the impact of the mating system. The length of the genomic region on which a second mutation can appear was chosen arbitrarily, and changing it can also change the probability. However, the important point for the deleterious-mutation mechanism to work is that there exists a size for regions flanking mating-type loci that allows both inversions to appear and mutations to accumulate, so that inversions can trap several deleterious mutations when suppressing recombination. The value *d* = 10^6^ chosen here allows to cover such flanking regions.

We computed our estimate of the probability of deleterious mutations accumulation for *r* = 0.001 (the two loci are close, strongly linked), *r* = 0.01, *r* = 0.1, and *r* = 0.5 (the two loci are distant, unlinked). We considered several values of selfing and intra-tetrad mating rates *f* and *p*_*in*_ in order to assess the impact of the mating system on the probability of deleterious mutation accumulation near a mating-type locus.

## 3 Results

### 3.1 Deleterious mutations are almost surely purged in the partial dominance case, and can escape purge in the overdominance case

#### Partial Dominance scenario

Under partial dominance, we find that the dominant eigenvalue *ρ* of the matrix *C* is always negative or null (see App. **??** and **??** for more details on the proof and computations). Previous theoretical results on branching processes state that, when *ρ <* 0, the probability that the deleterious mutation is purged from the population before it reaches a substantial frequency is one, and the mean time of purging is finite (see the Methods section). In particular, the probability of purging does not depend on the mating system (*ρ <* 0 for any value of intratetrad, intertetrad and outcrossing rates), nor on the recombination probability, selection and dominance coefficients. The only exceptions are when the deleterious mutation is neutral (*s* = 0) or behaves as neutral (*h* = 0 and *r* = 0, the mutation is neutral when heterozygous and completely linked to one mating-type allele), in which case the dominant eigenvalue is 0. The mutation is still purged from the population but previous theoretical results on branching processes state that this can take a much longer time compared to the case where *ρ <* 0, as the mean purging time would be infinite (see the Methods section).

Taking *w*_22_ = 1 − *hs* and *w*_32_ = 1 in the model of Ewens, 1967, to mirror our partial dominance scenario, the dominant eigenvalue becomes 1 − *hs*. It is always smaller than one, except when *h* = 0 or *s* = 0, *i.e*. when the mutation is neutral in the heterozygous state. Except in those cases, the mutation is purged from the population with probability one. We therefore find the same results as Ewens, 1967, and we extend these results in the case where mating is not random among gametes. In particular, the mutation being neutral in the heterozygous case (*h* = 0) is not sufficient to prevent the purging probability to be one when mating is not random: the mutation has to be completely linked to a permanently heterozygous locus (*h* = 0 and *r* = 0).

#### Overdominance scenario

Under overdominance, the dominant eigenvalue *ρ* can take positive or negative values. When *ρ* is positive, the probability that the mutation escapes purging and that the number of mutation-carriers increases exponentially fast is strictly positive. The general conditions on the parameters for *ρ* to be positive in our model are given in App. **??**, but they are difficult to interpret. Below, we describe a few simple cases in order to elucidate the role of each parameter, and then we complement the analysis with a numerical approach.

Similarly to the partial dominance case, the dominant eigenvalue is 0 when the mutation is neutral (*s*_3_ = 0, which implies *s*_4_ = 0 as well). The dynamics of the *b*-subpopulation (*i.e*. mutation-carriers) is then critical, which means that the mutant is purged with probability 1 but the mean purging time can be arbitrarily long (as the average extinction time of a critical branching process is infinite, see the Methods section).

When the mutation is not neutral (*s*_3_ ≠ 0) but with no disadvantage to *BB* homozygotes (*s*_4_ = 0), we prove that *ρ <* 0 (see App. **??**), which means that the dynamics of the *b* subpopulation is subcritical and that the mutant allele is purged with probability 1. This shows that the overdominant mutant allele is not maintained in the population when wild-type homozygotes are not disfavored compared to heterozygotes at the load locus. This corresponds to a completely recessive mutation, and is in agreement with the results for the partial dominance case with *h* = 0.

When the mutant allele is completely linked to a mating-type allele (*r* = 0), or under complete outcrossing (*f* = 0), the dominant eigenvalue is equal to *s*_4_, the selection coefficient for the fitness reduction of the *BB* wild-type homozygotes. The dynamics of the *b* subpopulation is then supercritical, which means that there is a non-zero probability that the mutant allele is not purged and, instead, reaches a significant number of carriers. Moreover, the mutant allele is more favored in this case when selection against *BB* homozygotes is stronger as it induces a stronger advantage of the *Bb* heterozygotes. A similar result can be derived from the work of Ewens, 1967. Taking *w*_22_ = 1 and *w*_32_ = 1 − *s*_4_ in his model to mirror our overdominance scenario, the dominant eigenvalue of Eq. **??** becomes 1/(1 − *s*_4_). As long as *s*_4_ > 0, this eigenvalue is always greater than one, and its value increases as the selection against wild-type homozygotes increases. This shows that the dynamics of an overdominant allele under random gamete mating is similar as under complete outcrossing.

In the case of complete intra-tetrad selfing (*f* = 1, *p*_*in*_ = 1), we find that *ρ* ≥ 0 if *r* ≤ 2*s*_4_, in agreement with the results of Antonovics and Abrams, 2004. These results mean that the overdominant mutation can be maintained under complete selfing if it is tightly linked to the mating-type locus (*r* small) or if the heterozygote advantage over wild-type homozygotes is strong (*s*_4_ large).

In the case of complete selfing (*f* = 1), we find that *ρ* = *s*_4_ − *s*_3_ ≤ 0 when *r*(2 − *r* − *p*_*in*_(1 − *r*)) − 2*s*_3_ ≥ 0. This shows that the dominant eigenvalue depends only on the selection coefficients when the recombination rate *r* exceeds a certain threshold (visible on the bottom panels of Figure **??**). This means that, if the recombination rate is larger than the strength of the selection against deleterious homozygotes, the mutation is purged with probability one. Moreover, the purging time is shorter when the difference in fitness between the two homozygotes is larger. The threshold on recombination increases as *p*_*in*_ increases, which means that the strength of the linkage between the mating-type locus and the mutation has the highest effect under intra-tetrad selfing.

Figure **??** shows more generally that the mating system affects the purging of deleterious mutations. On Figure **??**, the probability of purging is one in blue areas (the dominant eigenvalue is negative), and positive but smaller than one in red areas (the dominant eigenvalue is positive). The lines below which the mutation has a non-zero survival probability under the framework of Antonovics and Abrams, 2004, *i.e. r* = 2*s*_4_ under complete intra-tetrad selfing, are displayed as well. Comparing the panels for different values of intratetrad mating rate (*p*_*in*_) and selfing rate (*f*) shows that selfing favors the purging of the mutant allele (the blue area becomes larger as *f* increases), whereas intratetrad mating favors the maintenance of the deleterious allele (the blue area become smaller as *p*_*in*_ increases). Indeed, selfing favors the creation of homozygous individuals, which are disfavored, and intra-tetrad selfing favors the creation of heterozygous individuals, which are favored, compared to inter-tetrad selfing: the probability that a heterozygous individual *Bb* produces a heterozygous offspring *Bb* is higher under intra-tetrad selfing (probability 1 − *r*/2) than under inter-tetrad selfing (probability 1 − *r* + *r*^2^/2).

**Figure 2:**
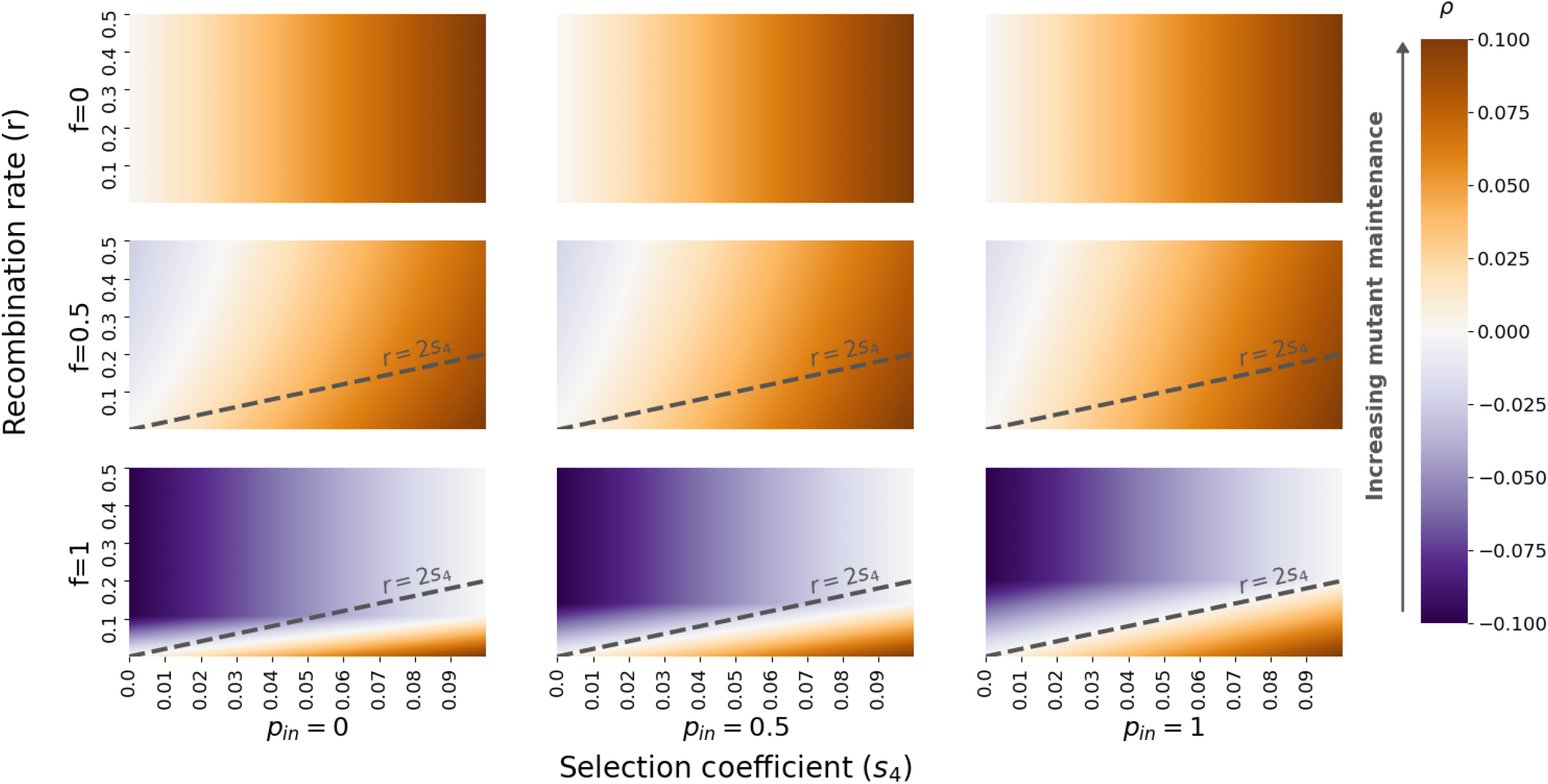
Dominant eigenvalue *ρ* for the overdominance scenario. When *ρ ≤* 0 (blue areas), the mutation is purged with probability 1. When *ρ* > 0 (red areas), the mutation has a non-zero probability to escape purging. The mutation is maintained longer in the population as *ρ* increases. All panels have the same axes. x-axis: *s*_4_, selection coefficient for wild-type *BB* homozygotes. y-axis: *r*, recombination rate between the two loci. Each column corresponds to a value of *p*_*in*_ (intra-tetrad rate, 0, 0.5, 1), and each row to a value of *f* (selfing rate, 0, 0.5, 1). The selection coefficient for *bb* homozygotes is set to *s*_3_ = 0.1. The line *r* = 2*s*_4_ is displayed for comparison with the findings in Antonovics and Abrams, 2004.

### 3.2 The presence of a mating-type locus has a sheltering effect under partial selfing

Looking at the derivative of the dominant eigenvalue at *r* = 0.5, we find that the presence of a mating-type locus near the mutation has a sheltering effect on the deleterious mutation, under partial selfing and in both selection scenarii. Indeed, the derivative 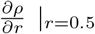 is always negative, except when the mutation is neutral (*s* = 0 under partial dominance or *s*_3_ = 0 under overdominance), when it is lethal (*s* = 1) or dominant (*h* = 1) under partial dominance, or under complete outcrossing (*f* = 0) in both scenarii, in which cases the derivative is zero and there is no sheltering effect. Under complete selfing (*f* = 1), the derivative is also null when the intratetrad coefficient *p*_*in*_ is below a certain threshold (see App **??** and **??** for the proof). As explained in the Methods section, this analysis shows that, in a wide range of situations, the rate of decay of the mutant subpopulation is lower when the mutation is linked to a mating-type locus, even loosely (*i.e*. as soon as *r <* 0.5), than when recombination is free between the two loci. Hence, except for the particular cases cited above, the mating-type locus always has a sheltering effect on the deleterious mutation maintenance under partial selfing, independently of the mating system coefficients (*f* and *p*_*in*_) and of the selection and dominance coefficients (*s* and *h*, or *s*_3_ and *s*_4_).

Figure **??** shows that, under both partial dominance or overdominance, the variation of the derivative at *r* = 0.5 is stronger when the selfing rate *f* (x-axis) or the intratetrad selfing probability *p*_*in*_ (y-axis) are high. This means that the sheltering effect of the mating-type locus is stronger under high selfing or high intratetrad mating. Two forces oppose here: increasing selfing induces a greater production of homozygotes, which are disfavored, whereas increasing intra-tetrad selfing rate or increasing the linkage with a mating-type locus favors the production of heterozygotes, which are favored. The sheltering effect of the mating-type locus that counters the purging effect of selfing is higher when selfing is higher, and this countering effect is reinforced by a high intra-tetrad mating rate. Moreover, when approaching *f* = 1, the derivative decreases to 0. Indeed, the selection and dominance coefficients *s, s*_*3*_ and *h* are here sufficiently small for the condition to have 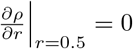 when *f* = 1 to be met, for both selection scenarii (see App. **??** and **??** for the derivation of this condition). This means that the dynamics of the deleterious mutation is independent of the presence of a mating-type locus under complete selfing and weak selection.

We explore the impact of other parameters in the Supplementary materials. Figure **??** shows that, under partial dominance, the sheltering effect of a mating-type locus is stronger when the dominance coefficient *h* is lower (*Bb* heterozygotes, which are more prone to be created in the presence of a mating-type locus, are more favored) or when the selection coefficient *s* is high (the differential in fitness between *Bb* heterozygotes and *bb* homozygotes is higher). Similarly, Figure **??** shows that, under overdominance, the sheltering effect of the mating-type locus is stronger when the selection against *bb* homozygotes is higher (*s*_3_ coefficient), whereas the selection against *BB* homozygotes does not impact the strength of the sheltering effect, suggesting that the dynamics of the deleterious allele is mostly driven by the difference in fitness between the favored heterozygotes and the disfavored deleterious homozygotes.

Looking at the derivative at *r* = 0, we show in App. **??** and App. **??** that it is also negative in both selection scenarii. This means that the eigenvalue decreases, *i.e*. that the mutation is less maintained in the population as soon as the two loci are no longer completely linked. Figure **??** shows that the difference 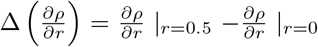 is always positive, which means that the absolute value of the derivative at *r* = 0 is larger than the absolute value of the derivative at *r* = 0.5. This shows that the sheltering effect is stronger on mutations closely linked to the mating-type locus : adding a small chance of recombination on previously completely linked loci (*r* = 0) has a greater impact on the maintenance of deleterious mutations than adding a small amount of linkage between two previously completely unliked loci (*r* = 0.5). The largest difference between the two derivatives occurs for selfing rates close to one, the derivative being then zero at *r* = 0.5, while the derivative at *r* = 0 approaches −1. This shows that the linkage to the mating-type locus particularly impacts the strength of its sheltering effect under high selfing.

**Figure 3:**
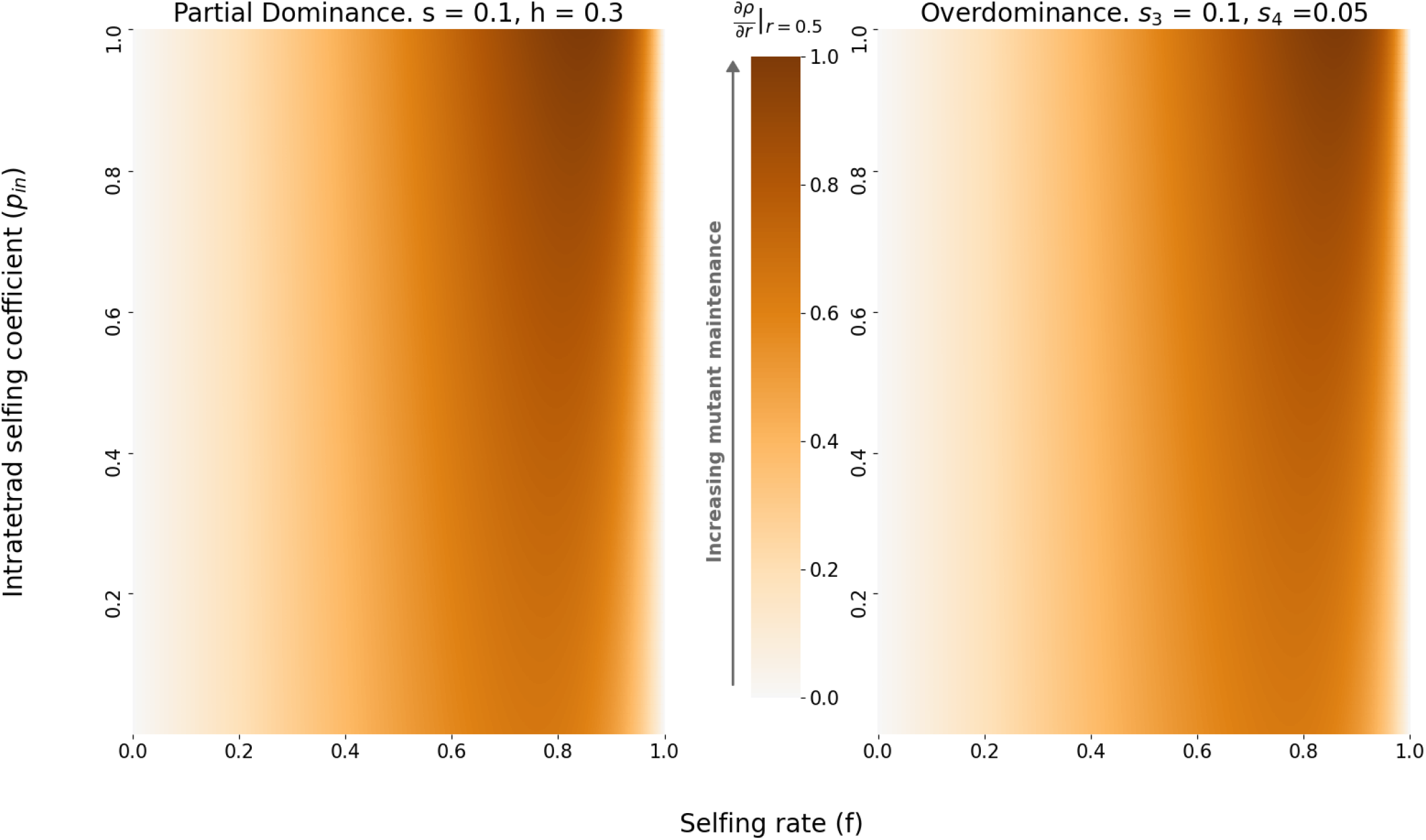
Relative variation of the derivative of the dominant eigenvalue in the partial dominance case (left) and the over-dominance case (right). For each panel, the values 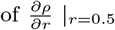 range from a minimal value, which is negative, to zero. We divided each value of the derivative by this minimum in order to plot values between 0 and 1 for every panel. This enables us to compare the effect of the presence of a mating-type locus on the same scale for both selection scenarii. x-axis: selfing rate *f*. y-axis: intratetrad selfing rate *p*_*in*_. The darker the color, the more the mating-type locus shelters the mutation, thus promoting its maintenance.

### 3.3 Rare events of maintenance of the deleterious mutation occur in both selection scenarii, paving the way for an accumulation of mutations

The empirical distribution of the purging time of the deleterious mutation in the partial dominance case is shown on Figure **??**: for ca. 75% of the independent runs, the mutation was rapidly purged, while in some rare cases (ca. 1%), the purge took very long (several orders of magnitude longer than the 75% percentile empirically obtained from the 100,000 runs). Note that the approximation of the distribution of the time to extinction by a Gumbel law (Th. 4.1 of Heinzmann, 2009) falls short here, because the initial number of individuals (one) and the absolute value of *ρ* (given in the caption) are too small.

Consistently with our results that 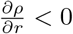, the sheltering effect of the mating-type locus implies that the purging time increases when the recombination rate decreases (Figure **??**, and Figure **??** for the overdominant case). We also consistently find that increasing selfing decreases the purging time (Figures **??** and **??**). In each case, the closer *ρ* is to zero, the more extreme the rare events are : the distribution of the 1% longest purging times is stretched towards higher values when *ρ* gets closer to zero, while the distributions of the 75% shortest remain similar.

Figure **??** displays the probability that the mutation can be maintained long enough in the population for another mutation to appear in a region of 10^6^ bp near the mating-type locus. This probability is nonnegligible (of the order of 1% to 10%), which shows that accumulation events are rare but still occur near mating-type loci. This is true even under selfing as the sheltering effect of the mating-type locus can counter the purging effect of selfing. Indeed, when the recombination rate between the first mutation and the mating-type locus is high (*r* = 0.5 or *r* = 0.1), modeling a situation where the distance between the two loci is large, the probability that a second mutation appears before the first one is purged decreases with increasing selfing, even with high intra-tetrad selfing rates. However, when the first mutation is closer to the mating-type locus (lower recombination rates), the probability that a second mutation appears before the first one is purged under selfing is similar to the probability under complete outcrossing. The presence of a mating-type locus can thus facilitate the accumulation of deleterious mutations in its flanking regions, especially in highly selfing populations.

**Figure 4:**
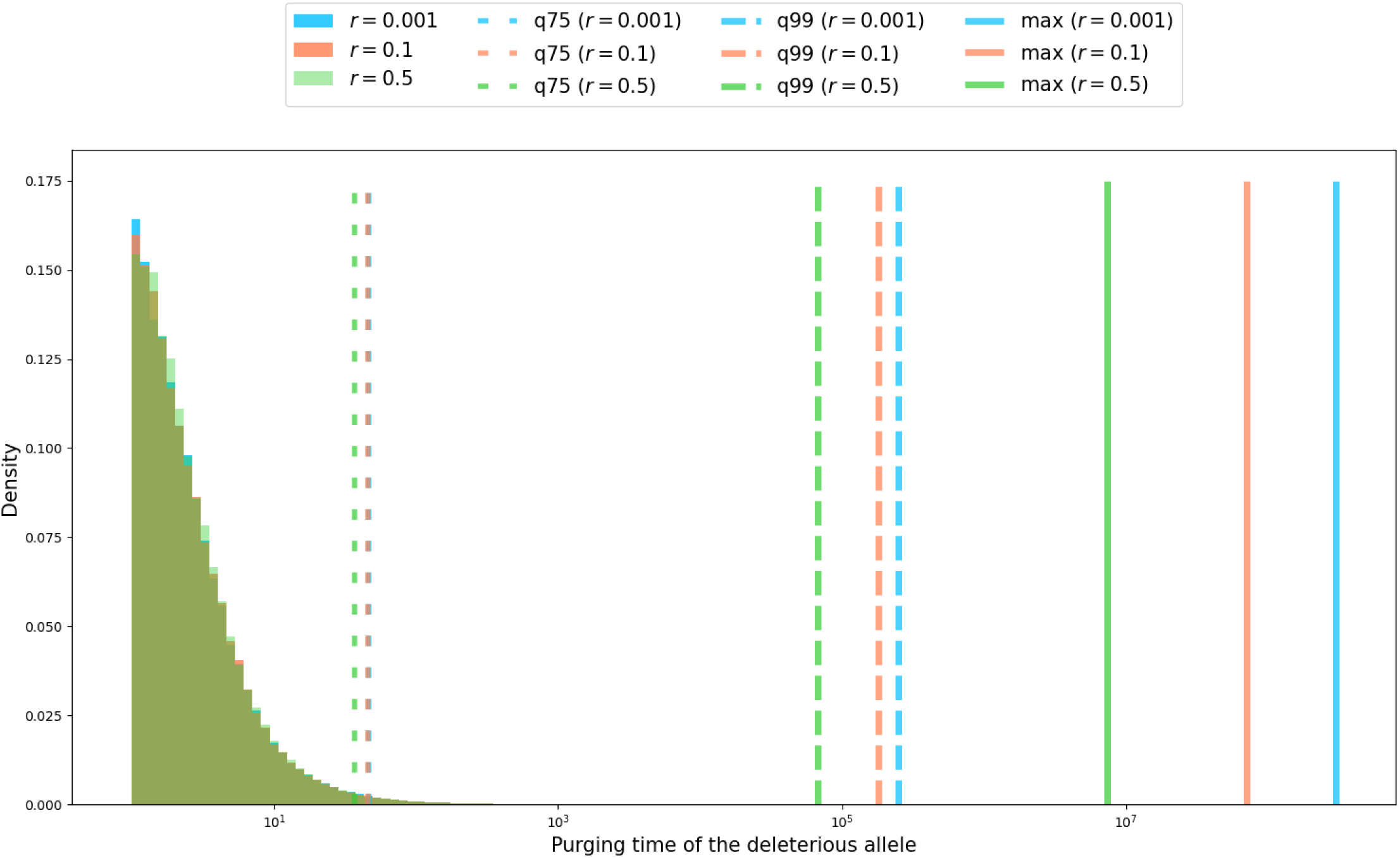
Empirical distribution of the deleterious allele purging time for the partial dominance scenario. A total of 100,000 simulations were run, with *s* = 0.1, *h* = 0.1, *f* = 0.5, *p*_*in*_ = 0.5, starting from one heterozygous individual (*X*_0_ = (1, 0, 0)), and for three values of the recombination rate (*r* = 0.001 in blue, *r* = 0.1 in red and *r* = 0.5 in green). The respective values for *ρ* are *ρ* = − 0.0101, *ρ* = −0.0106 and *ρ* = − 0.0307. The x-axis is log-scaled. The large-dotted lines represent the 75^*th*^ percentile (*q75*), the dashed lines indicate the 99^*th*^ percentile (*q99*), and solid lines the maximum value (*max*) of the purging time. Maximum values are several order of magnitudes higher than the 75th percentile of the empirical distribution of the purging time.

**Figure 5:**
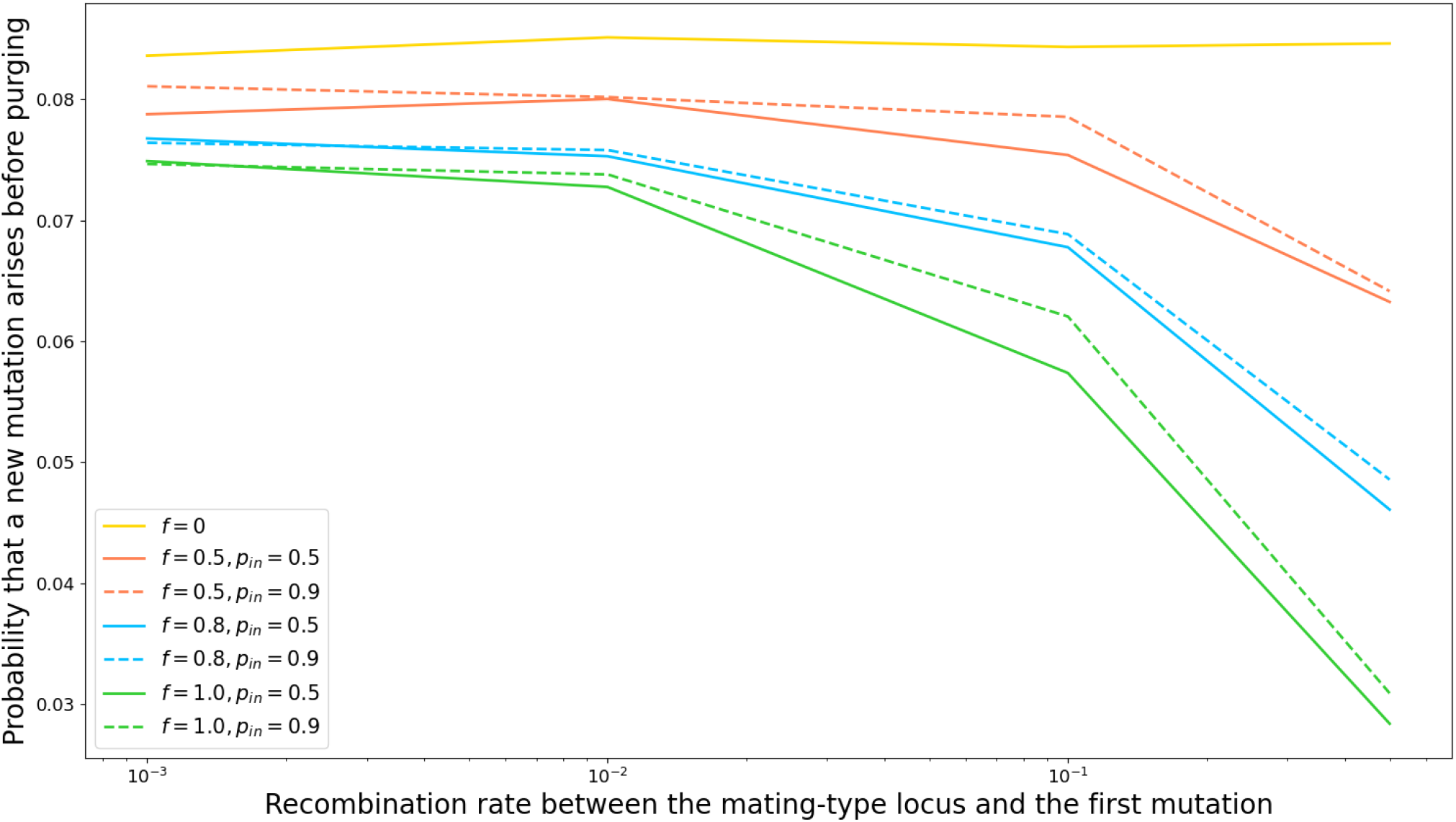
Probability that a new mutation appears in a region of length 10^6^ bp before the first mutation is purged from the population, under the partial dominance scenario, depending on the recombination rate between the first mutation and the mating-type locus. We considered a mutation rate per base pair per reproduction event of 10^−8^. Here, the reproduction events are those of the branching process, that change the composition of the mutant-carriers subpopulation. The probability that a new mutation appears before the purge of the first one is approximated by the proportion of simulation runs for which the number of reproduction events exceeds the expected number of events needed for a new mutation to appear (see text). For each set of parameters (*r, f, p*_*in*_), 100,000 independent simulations were run. Colors correspond to different values of the selfing rate *f*, and line styles to different values of the intra-tetrad selfing rate *p*_*in*_. When *f* = 0, a single curve is displayed, as the value of *p*_*in*_ has no impact under complete outcrossing. For all simulations, we set *s* = 0.1 and *h* = 0.1.

## 4 Discussion

### Partially recessive deleterious mutations are almost surely purged in finite time while overdominant mutations can persist

We have shown that partially recessive deleterious mutations close to a fungal-like mating-type locus (*i.e*. that does not prevent diploid selfing) are almost surely purged in finite time, except when they are neutral or behave as neutral. In the overdominance case, the probability of purge depends on parameter values. Low selfing rates, high intra-tetrad selfing rates or tight linkage to the mating-type locus increases both the maintenance probability and persistence of the overdominant allele, whereas a high selfing rate favors its purge.

In particular, if linkage is complete (corresponding to *r* = 0 here, or to the case where the inversion encompasses a permanently heterozygous locus in Jay et al., 2022), an overdominant allele may be maintained in a population and even sweep to fixation with non-zero probability, which confirms previous findings (Antonovics et al., 1998, Antonovics and Abrams, 2004, Jay et al., 2022). This means that, although selfing purges deleterious mutations, a mating-type locus can have a sheltering effect in its flanking regions.

In general, the overdominant allele is maintained longer and with a higher probability in the population when the fitness advantage of heterozygotes over homozygotes is higher, in line with previous simulation results (Antonovics and Abrams, 2004). This conclusion is sensible: if the mutant is strongly favored in a heterozygous state, it can be maintained in this state in the population.

### The presence of the mating-type locus has a sheltering effect under selfing

We found that, in both selection scenarii, the presence of the mating-type locus had no effect on the maintenance of deleterious mutations under outcrossing, but always had a sheltering effect under selfing, which strengthened as the selfing rate increased. Indeed, selfing increases homozygosity and thus accelerates the purge of a deleterious allele, whereas the presence of a permanently heterozygous mating-type locus induces more heterozygosity in its flanking regions, that counters the purging effect of selfing. The sheltering effect of a mating-type locus is thus all the more tangible as it counters the strong purging effect induced by selfing. Increasing intra-tetrad selfing also induces more heterozygosity and thus slightly reinforces the sheltering effect of the mating-type locus. This is consistent with the findings that, in fungi, ascomycetes that reproduce via outcrossing and live as haploids do not show evolutionary strata (Skinner et al., 1993, Zhong et al., 2002, Phan et al., 2003, Kuhn et al., 2006, Jin et al., 2007, Malkus et al., 2009) whereas pseudo-homothallic ascomycete fungi, living as dikaryotic and undergoing mostly intra-tetrad selfing, are those with evolutionary strata around their mating-type locus (Menkis et al., 2008, Hartmann, Duhamel, et al., 2021, Hartmann, Ament-Velásquez, et al., 2021, Vittorelli et al., 2022). In basidiomycetes also, the species with evolutionary strata are dikaryotic and automictic, *e.g. Microbotryum fungi* and *Agaricus bisporus* var. *bisporus* (Branco et al., 2017, Branco et al., 2018, Foulongne-Oriol et al., 2021). This may be explained by the fact that intra-tetrad selfing favors the accumulation of deleterious alleles near the mating-type locus, which in turn can promote selection for recombination suppression because there will be more variability in the number of mutations present in a genomic region close to the mating-type locus, and therefore more fragments having a much lower number of deleterious mutations than average in the population (Jay et al., 2022).

Additionally, we found that the sheltering effect of a mating-type locus was stronger when the mutation was more strongly recessive. Indeed, the purging effect of selfing on partially recessive mutations is stronger for more recessive mutations (Charlesworth and Charlesworth, 1987, Caballero and Hill, 1992, Arunkumar et al., 2015), in which case the opposite force of the sheltering effect of a mating-type locus is strenghtened. This is in agreement with the results of studies on the sheltered load linked to a self-incompatibility locus, showing that completely recessive deleterious mutations are more easily fixed than partially recessive ones (Llaurens et al., 2009). This also confirms results on the fixation of inversions encompassing recessive deleterious mutations and linked to a permanently heterozygous locus (Olito et al., 2022, Jay et al., 2022). These results showed that inversions became fixed with a higher probability when segregating deleterious mutations were more strongly recessive.

### Rare events of long maintenance of deleterious mutations in the population can occur

We further found that rare events of long maintenance of deleterious mutations in the population occurred under both selection scenarii. This shows that some deleterious mutations can persist in the population for an extended period of time before being purged, especially near the mating-type locus: in approximately 1% of our simulations, the purge of the deleterious mutation took several orders of magnitude longer than the 75% percentile empirically obtained from the 100,000. These surprisingly long purging times are likely to be due to the dynamics of the mutant being almost critical (the dominant eigenvalue in the branching process approximation is negative, but close to zero). However, from a modeling perspective very little is currently known about these trajectories, and more generally about the extinction time of multitype branching processes. Studying the extinction time of a deleterious allele in a one locus-two allele setting with a unitype branching process approximation and a diffusion approximation showed that the standard deviation of the mean extinction time was higher than the mean itself (Nei, 1971), which is a feature that was also found in our simulations of multitype branching processes. These results show that the extinction time of deleterious alleles is highly variable, producing long-lasting mutations that may induce an accumulation of deleterious alleles near a mating-type locus, which is a prerequisite for recombination suppression to extend away from this locus (Jay et al., 2022).

### The dynamics of deleterious mutations heavily relies on the mating system

Our results show that the mating system, and selfing in particular, is a prevailing force impacting the dynamics of deleterious mutations. Indeed, we found that a mating-type locus shelters mutations and thus favors their maintenance, but increasing selfing reduces the maintenance of mutations with a stronger effect. This result is congruent with previous studies showing that an increase in the selfing rate induces i) a reduction of the mutational load at a given locus or at multiple non-interacting loci far from mating-type compatibility loci (Charlesworth et al., 1990, for a deterministic model, Abu Awad and Roze, 2018, for diffusion approximation), and ii) a reduction of the purging time of deleterious mutations (Caballero and Hill, 1992).

However, we observed a particular behavior when the population reproduced only via selfing. Under complete selfing in our setting, the existence of a sheltering effect of a mating-type locus strongly depended on the values of the intra-tetrad selfing rate: the sheltering effect of the mating-type locus was detectable only when the intra-tetrad selfing coefficient exceeded a certain threshold, that depended on the dominance and selection coefficients. This strong effect of departing from complete selfing had previously been noted: introducing a small amount of outcrossing in a selfing population can lead to sharp changes in the dynamics of a deleterious mutation, whereas adding a small amount of selfing in an outcrossing population induces a smoother change (Holsinger and Feldman, 1985).

### Limits of the methods

Our results are limited to the case of a single load locus, in interaction with a heterozygous mating-type locus, and may not apply when considering different frameworks, such as multiple epistatic loci or with additional beneficial mutations, especially regarding the impact of the mating system. Indeed, selfing has a non-monotonous effect depending on the tightness of linkage between multiple interacting loci (Abu Awad and Roze, 2018): at low selfing rates, increasing linkage between loci increases the mutation load, whereas the opposite effect is observed at high selfing rates. Selfing also has a non-monotonous effect on genetic variation in populations under stabilizing selection (Lande and Porcher, 2015, Clo and Opedal, 2021). In addition, selfing can enhance the fixation chances of a deleterious allele when it hitchhikes during a selective sweep (Hartfield and Otto, 2011, Hartfield and Glémin, 2014). Moreover, the impact of the mating system on the maintenance of deleterious mutations may be different if the number of individuals carrying the mutant allele exceeds a certain threshold. In this case, the branching process approximation does not hold anymore, and a deterministic model in large population may be used to further describe the dynamics of the deleterious allele (Durrett and Schweinsberg, 2004, Durrett, 2008 Section 6.1.3). The impact of the mating system then remains unclear: in large populations, selfing reduces the effective population size, which impairs the efficiency of selection and increases the mutational load of the population, but it also bolsters homozygosity, which favors the purge of deleterious mutations (Pollak, 1987, Caballero and Hill, 1992, Charlesworth and Wright, 2001, Wright et al., 2008).

Another limitation of our approach is that we considered a fixed recombination rate for simplicity, but allowing this rate to vary would allow us to test whether recombination suppression could evolve. Such an outcome may depend on the strength of selection against the deleterious mutation, as well as on the mating system (Antonovics and Abrams, 2004, Abu Awad and Roze, 2018). In some previous models, the impact of a modifier of recombination in the form of a multi-allelic locus was studied by simulations, but no analytical results were obtained (Feldman, 1972, Palsson, 2002, Antonovics and Abrams, 2004, Lenormand and Roze, 2022). The multitype branching process framework developed here would also be an interesting approach to obtain numerical results on this more complex situation, but analytical results would probably be out of reach because of the increase in complexity of the model.

### Conclusion and Perspectives

In conclusion, our findings show that a mating-type locus has a sheltering effect on nearby deleterious mutations, especially in case of selfing and automixis, which can then play a role in the evolution of recombination suppression near mating-compatibility loci (Antonovics and Abrams, 2004, Jay et al., 2022). This may contribute to explain why evolutionary strata of recombination suppression near the mating-type locus are found mostly in automictic (pseudo-homothalic) fungi (Menkis et al., 2008, Branco et al., 2017, Branco et al., 2018, Hartmann et al., 2020, Hartmann, Ament-Velásquez, et al., 2021, Foulongne-Oriol et al., 2021, Vittorelli et al., 2022).

The results obtained here on the accumulation of deleterious mutations should apply, beyond fungal-like mating-type loci, to other permanently heterozygous loci, such as supergenes (Llaurens et al., 2017). In contrast, sporophytic or gametophytic plant self-incompatibility loci prevent diploid selfing, leading to a completely different evolutionary scenario in their flanking regions as imposed by complete outcrossing. The diversity of observed patterns regarding the presence or absence, length and number of evolutionary strata around these regions (Uyenoyama, 2005) may be explained, in addition to the mating system, by other factors controlling the long-term behavior of deleterious mutations which are not studied here, such as the number of alleles at supergenes, the length of the haploid phase (Jay et al., 2022), or the presence of multiple load loci that are possibly physically linked and with epistatic interactions (Abu Awad and Roze, 2018, Lenormand and Roze, 2022). The questions of the genome-wide impact of a mating-type locus, and of the interaction between a permanently heterozygous locus and background mutations, are currently debated (Abu Awad and Waller, 2021). The branching process framework developed here could be applied to diploid individuals carrying a load locus with two alleles, undergoing selfing or outcrossing, in order to investigate the dynamics of a new deleterious mutation in a population with or without a mating-type locus.

Our results showing the long maintenance of deleterious mutations in the vicinity of permanently heterozygous loci pave the way for future investigations on the accumulation of deleterious mutations. Previous studies (Coron et al., 2013, Coron, 2014) on mutational meltdown, showing that deleterious mutations accumulate faster when other mutations are already fixed, also encourage future work in this direction.

## Acknowledgments

We thank Paul Jay and Denis Roze for insightful discussions, Aurélien Tellier for handling the recommendation process, and three anonymous reviewers for their useful and very constructive comments.

## 5 Appendix: Table of notation

**Table 1.**

|  |  |
| --- | --- |
| $N$ | Population size |
| $G_1, \dots, G_4$ | Genotypes |
| $(g_1, \dots, g_4)$ | Number of individuals of each genotype |
| $f$ | Selfing probability |
| $p_{in}$ and $p_{out} = 1 - p_{in}$ | Intra- and Inter-tetrad selfing probabilities |
| $r$ | Recombination rate |
| $S_i$ | Probability of survival of an offspring of genotype $i \in \{1, 2, 3, 4\}$ (see Figure ??) |
| $s$ | Selection coefficient in the partial dominance case |
| $h$ | Dominance coefficient in the partial dominance case |
| $s_3, s_4$ | Selection coefficients in the overdominance case |

## 6 Appendix: Intra-, Inter-tetrad selfing and outcrossing

**Figure 6:**
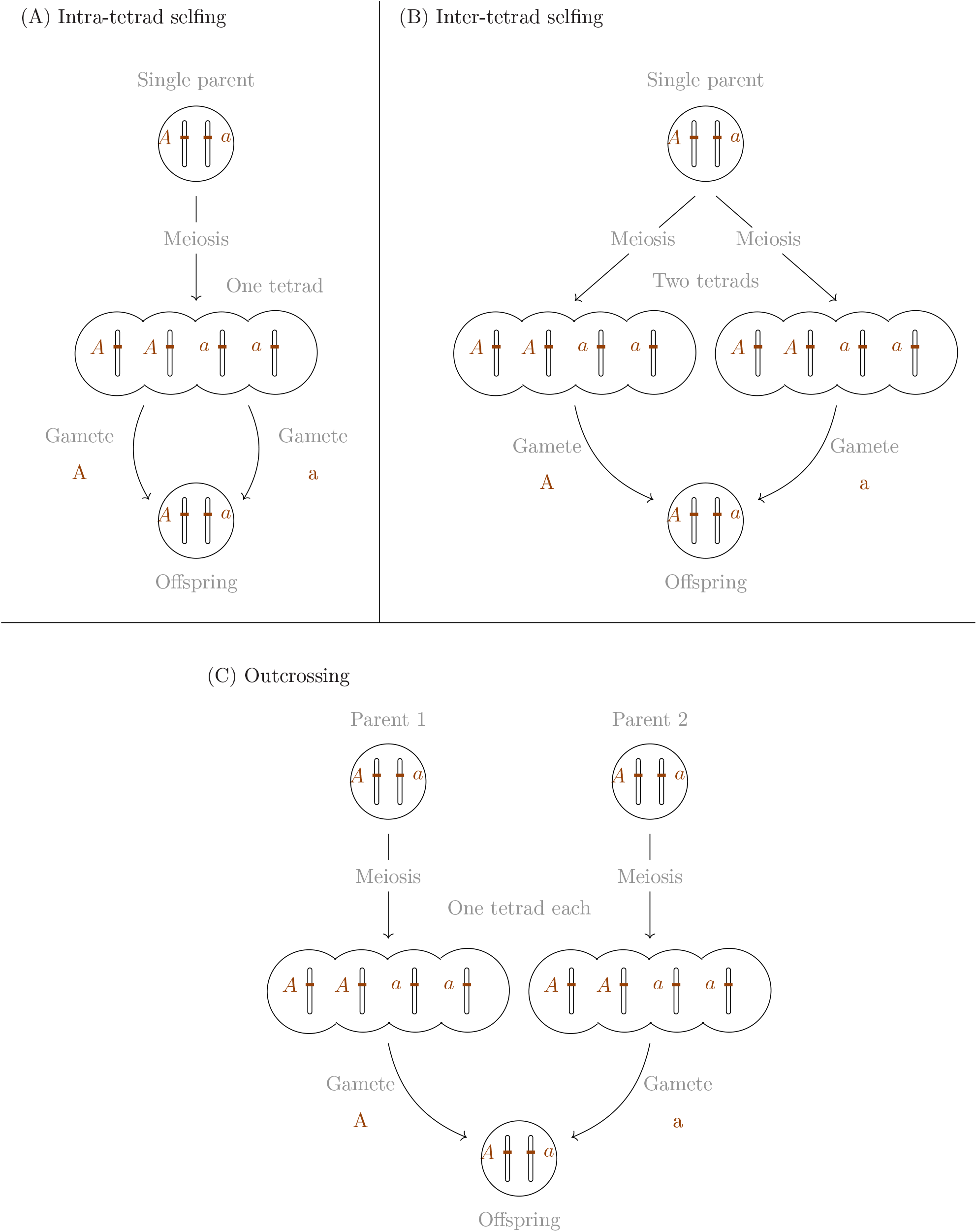
Schematic representation of the three mating systems considered in the model. Individuals are represented by a pair of mating-type chromosomes, with the mating-type locus displayed. A diploid offspring is generated by the fusion of two gametes carrying different mating-type alleles (A and a). (A) Under intra-tetrad selfing, both gametes are picked from the same tetrad; only one parent is involved. (B) Under inter-tetrad selfing, the two gametes are picked from two different tetrads (meioses) produced by the same diploid parent; only one parent is involved. (C) Under outcrossing, the two gametes are picked in tetrads produced by different parents.

## 7 Appendix: Appendices for the Method section

### 7.1 Rates of creation of offspring with given genotypes (Moran process)

**Table 2:**
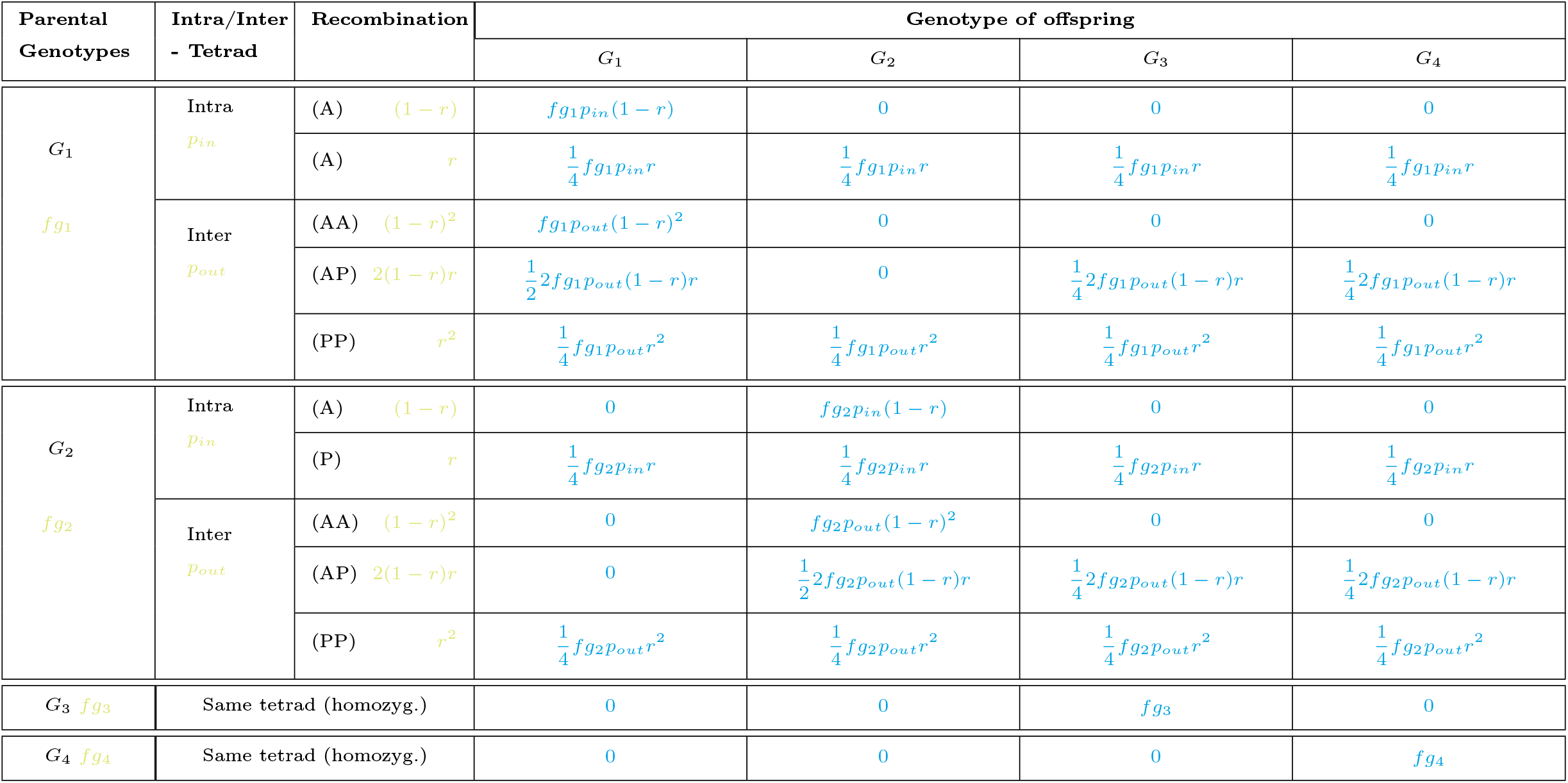
Table summarizing the rates of production of an offspring of each genotype (last four columns) in case of **selfing**. *Parental Genotype*: The genotype of the individual involved in the mating event; *Intra/Inter-tetrad* : Mating through intra-of inter-tetrad selfing (see section **??** for definitions); *Recombination*: Occurrence of a recombination event in the tetrads from which gametes are picked. “A” stands for “Absence” in one tetrad, “P” stands for “Presence” in one tetrad. We use only one letter when the two gametes come from the same tetrad or when one of the genotypes involved is homozygous at the load locus. For example, (AP) indicates that recombination occured in one tetrad but not in the other. *G*_*i*_: the rate at which an offspring of genotype *G*_*i*_ is produced, due to the scenario of parental genotype, intra/inter tetrad selfing and presence/absence of recombination considered. The total rate *T*_*g*_ (+*G*_*i*_) at which a new offspring of genotype *G*_*i*_ is created when the population state is *g* = (*g*_1_, *g*_2_, *g*_3_, *g*_4_) is then the sum of all the rates appearing in column *G*_*i*_ in this Table, Table **??** and Table **??**.

| Parental Genotypes | Intra/Inter - Tetrad | Recombination | Genotype of offspring |  |  |  |
| --- | --- | --- | --- | --- | --- | --- |
| | | | $G_1$ | $G_2$ | $G_3$ | $G_4$ |
| $G_1$<br>$fg_1$ | Intra<br>$p_{in}$ | (A) $(1-r)$ | $fg_1p_{in}(1-r)$ | 0 | 0 | 0 |
| | | (A) $r$ | $\frac{1}{4}fg_1p_{in}r$ | $\frac{1}{4}fg_1p_{in}r$ | $\frac{1}{4}fg_1p_{in}r$ | $\frac{1}{4}fg_1p_{in}r$ |
| | Inter<br>$p_{out}$ | (AA) $(1-r)^2$ | $fg_1p_{out}(1-r)^2$ | 0 | 0 | 0 |
| | | (AP) $2(1-r)r$ | $\frac{1}{2}2fg_1p_{out}(1-r)r$ | 0 | $\frac{1}{4}2fg_1p_{out}(1-r)r$ | $\frac{1}{4}2fg_1p_{out}(1-r)r$ |
| | | (PP) $r^2$ | $\frac{1}{4}fg_1p_{out}r^2$ | $\frac{1}{4}fg_1p_{out}r^2$ | $\frac{1}{4}fg_1p_{out}r^2$ | $\frac{1}{4}fg_1p_{out}r^2$ |
| $G_2$<br>$fg_2$ | Intra<br>$p_{in}$ | (A) $(1-r)$ | 0 | $fg_2p_{in}(1-r)$ | 0 | 0 |
| | | (P) $r$ | $\frac{1}{4}fg_2p_{in}r$ | $\frac{1}{4}fg_2p_{in}r$ | $\frac{1}{4}fg_2p_{in}r$ | $\frac{1}{4}fg_2p_{in}r$ |
| | Inter<br>$p_{out}$ | (AA) $(1-r)^2$ | 0 | $fg_2p_{out}(1-r)^2$ | 0 | 0 |
| | | (AP) $2(1-r)r$ | 0 | $\frac{1}{2}2fg_2p_{out}(1-r)r$ | $\frac{1}{4}2fg_2p_{out}(1-r)r$ | $\frac{1}{4}2fg_2p_{out}(1-r)r$ |
| | | (PP) $r^2$ | $\frac{1}{4}fg_2p_{out}r^2$ | $\frac{1}{4}fg_2p_{out}r^2$ | $\frac{1}{4}fg_2p_{out}r^2$ | $\frac{1}{4}fg_2p_{out}r^2$ |
| $G_3$ $fg_3$ | Same tetrad (homozyg.) | | 0 | 0 | $fg_3$ | 0 |
| $G_4$ $fg_4$ | Same tetrad (homozyg.) | | 0 | 0 | 0 | $fg_4$ |

**Table 3:**
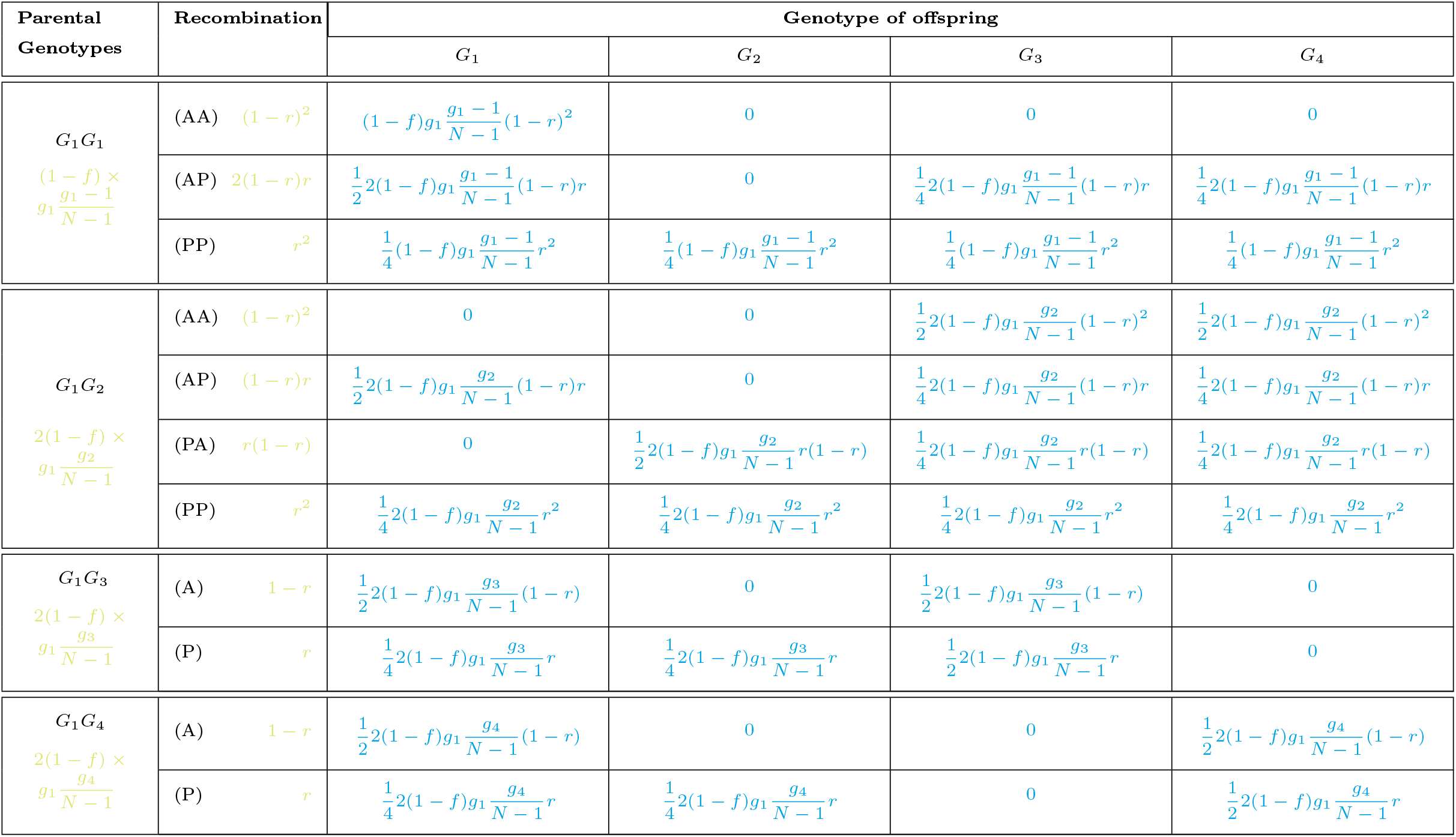
Part 1 of the table summarizing the rates of production of an offspring of each genotype (last four columns) in case of **outcrossing**. *Parental Genotype*: The genotype of the individuals involved in the mating event; *Recombination*: Occurrence of a recombination event in the tetrads from which gametes are picked. “A” stands for “Absence” in one tetrad, “P” stands for “Presence” in one tetrad. We use only one letter when the two gametes come from the same tetrad or when one of the genotypes involved is homozygous at the load locus. For example, (AP) indicates that recombination occured in one tetrad but not the other. *G*_*i*_: the rate at which an offspring of genotype *G*_*i*_ is produced, due to the scenario of parental genotype, intra/inter tetrad selfing and presence/absence of recombination considered. The total rate *T*_*g*_ (+*G*_*i*_) at which a new offspring of genotype *G*_*i*_ is created when the population state is *g* = (*g*_1_, *g*_2_, *g*_3_, *g*_4_) is then the sum of all the rates appearing in column *G*_*i*_ in this Table, Table **??** and Table **??**.

**Table 4:**
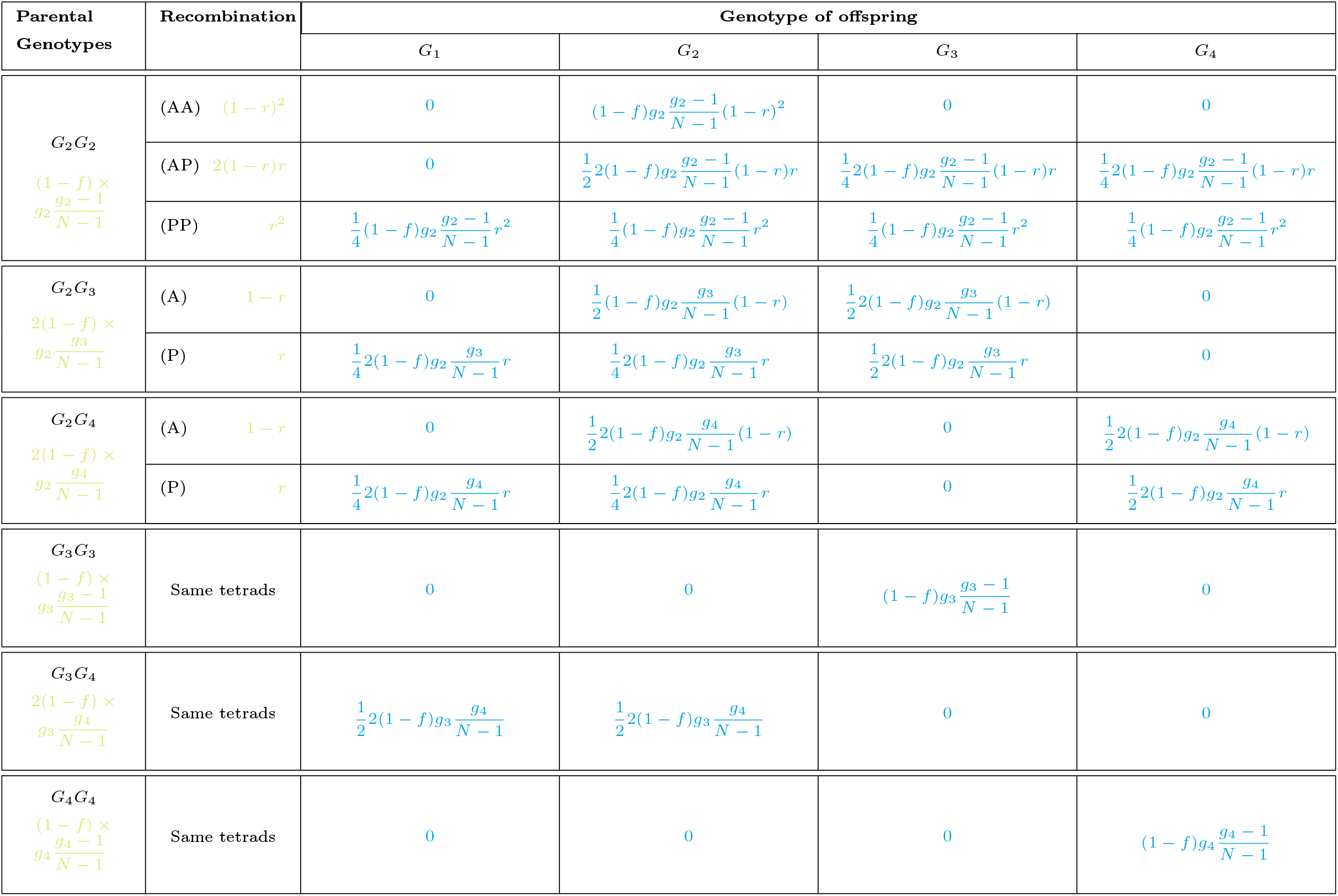
Part 2 of the table summarizing the rates of production of an offspring of each genotype (last four columns) in case of **outcrossing**. *Parental Genotype*: The genotype of the individuals involved in the mating; *Recombination*: Occurrence of a recombination event in the tetrads from which gametes are picked. “A” stands for “Absence” in one tetrad, “P” stands for “Presence” in one tetrad. We use only one letter when the two gametes come from the same tetrad or when one of the genotypes involved is homozygous at the load locus. For example, (AP) indicates that recombination occured in one tetrad but not the other. *G*_*i*_: the rate at which an offspring of genotype *G*_*i*_ is produced, due to the scenario of parental genotype, intra/inter tetrad selfing and presence/absence of recombination considered. The total rate *T*_*g*_ (+*G*_*i*_) at which a new offspring of genotype *G*_*i*_ is created when the population state is *g* = (*g*_1_, *g*_2_, *g*_3_, *g*_4_) is then the sum of all the rates appearing in column *G*_*i*_ in this Table, Table **??** and Table **??**.

The total rate at which an offspring of a given genotype is produced is then obtained by summing the rates along each column *G*_*i*_ in Tables **??, ??** and **??**. This gives:

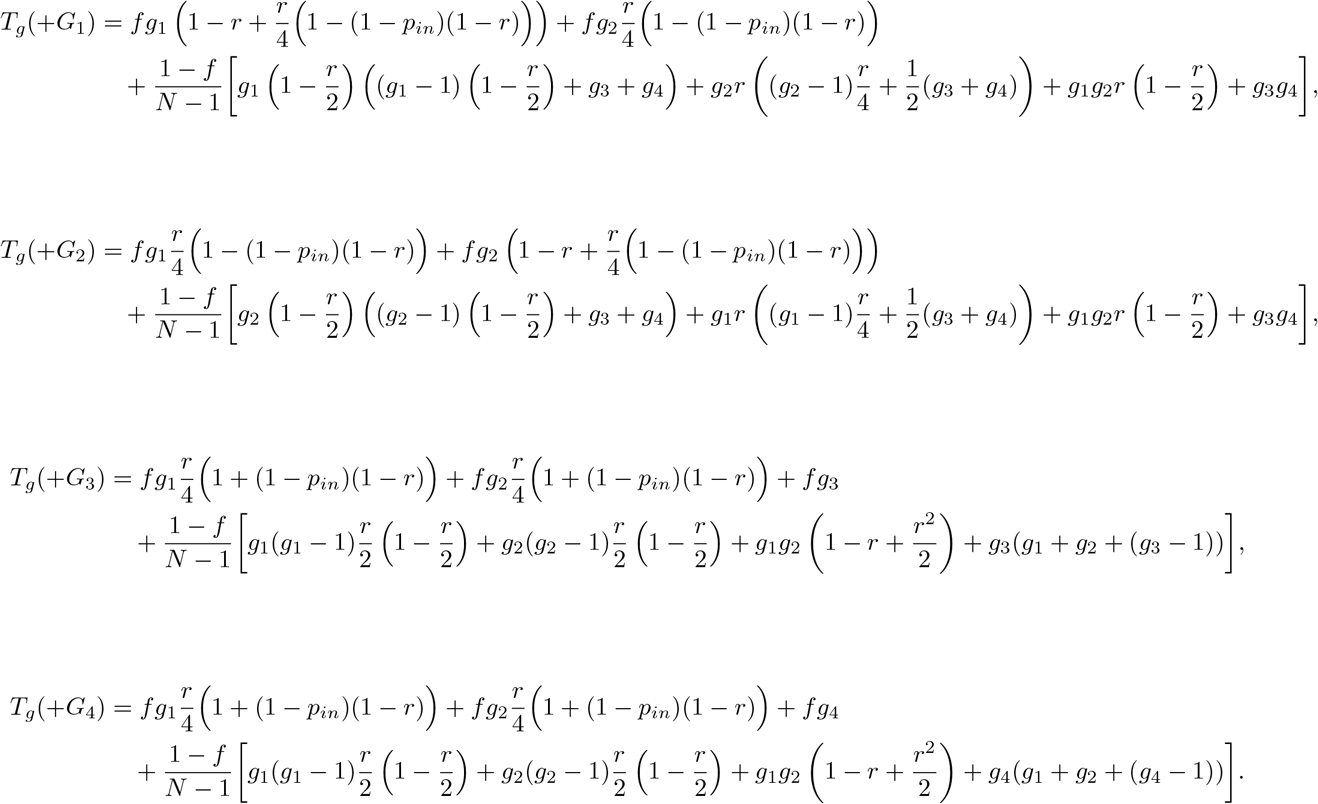

#### 7.2 Reproduction law for the branching process

We give here an example of how the reproduction laws for the branching process are derived from the rates of the Moran process, using the approximate regime (**??**).

Let us derive the coefficient *A*_12_ of the matrix *A*, which is the rate at which an individual of genotype *G*_2_ generates an offspring of genotype *G*_1_ and survives. Equivalently, this is the rate at which an individual of genotype *G*_2_ generates a descendance vector equal to *e*_1_ + *e*_2_.

Using the rates obtained for the Moran model, the rate at which an individual of genotype *G*_2_ produces an offspring of genotype *G*_1_ is:

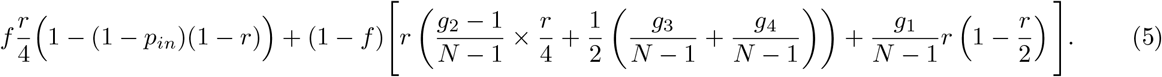

The first term, with a factor *f*, is the rate at which an individual of genotype *G*_2_ produces an offspring of genotype *G*_1_ by selfing. The second term, with a factor 1 − *f*, is the rate at which an individual of genotype *G*_2_ produces an offspring of genotype *G*_1_ by outcrossing. In this term, the fractions of the form 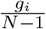 represent the probabilities that an individual of genotype *G*_*i*_ is chosen to mate with the *G*_2_ parent.

Using the approximation (**??**), *i.e*. assuming that *g*_4_ ≈ *N* and *g*_*i*_ « *N* for *i* = 1, 2, 3, we obtain that the quantity in Eq. (**??**) can be approximated by:

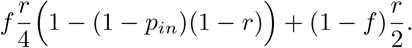

To obtain *A*_12_, it remains to multiply this rate by the probability that the offspring survives, *S*_1_, and the probability that the parent *G*_2_ is not chosen to die, 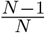. As the population size *N* is considered large, the latter probability is approximately equal to 1.

This gives:

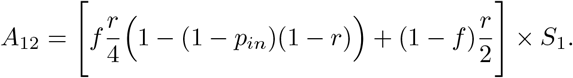

#### 7.3 Equation for the expected value of the size of the mutant population

This appendix gives the details of the derivation of the coefficients of the matrix *C* defined by

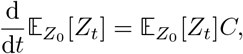

following Bacaër, 2018. Note that this is the same matrix defined in Athreya and Ney, 1972, Eq. 9, part. V.7.2., or in Pénisson, 2010, Eq. 1.1.16, but here we use the methodology described by Bacaër, 2018 to derive its coefficients.

In the following, type *j* refers to the genotype *G*_*j*_. We will use the standard notation 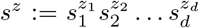 for *s* and *z* two vectors of the same dimension *d*.

##### 7.3.1 Notation

For all *t* ≥ 0, let us denote the expected value of the process at time *t* by *E*(*t*):

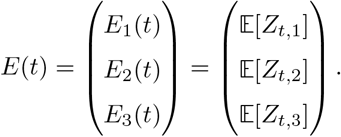

For *z* ∈ ℕ ^3^ and *t* ≥ 0, we let *p*(*t, z*) = ℙ(*Z*_*t*_ = *z*) be the probability that the system is found in state *z* at time *t*. Let *f* (*t*, .) be the generating function of the variable *Z*_*t*_: for all *s* ∈ [0, 1]^3^,

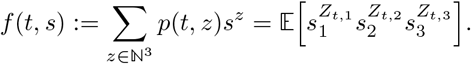

Recalling that *Y* ^*j*^ stands for the random vector of number of descendants of each type generated by the reproduction of a type *j* individual, we also define *π*_*j*_(*z*) = ℙ(*Y* ^*j*^ = (*z*_1_, *z*_2_, *z*_3_)). As indicated in the main text, the rates at which an individual of type *j* reproduces and gives rise to a descendance vector *e*_*i*_ + *e*_*j*_, *e*_*i*_ or 0 are respectively *A*_*ij*_, *T*_*ij*_ and *D*_*jj*_. We denote the total rate at which a reproduction event occurs for a parent of type *j* by 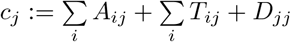.

The reproduction law of type *j* individuals is then given by, for every *i* ∈ {1, 2, 3},

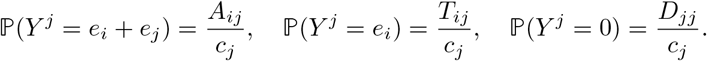

Finally, let *h*_*j*_ be the generating function of the reproduction law of type *j* individuals, for *j* ∈ {1, 2, 3}. That is, for *s* ∈ [0, 1]^3^,

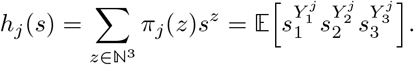

##### 7.3.2 Ordinary differential edecompoquation (ODE) satisfied by (*E*(*t*))_*t≥*0_

The reproduction law of each type has finite moments of all order, because the number of descendants produced can not exceed 2. That garantees that there is no explosion of the population in finite time. Hence, standard results on multi-dimensional random variables (see for example Athreya and Ney, 1972) give us that, for all types *j* and all *t* ≥ 0,

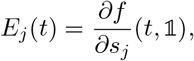

With 𝟙 = (1; 1; 1),, which gives

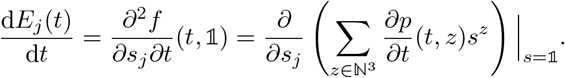

The variation of *p* over time 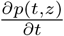 can be decomposed into two terms. For *z* ∈ ℕ^3^,

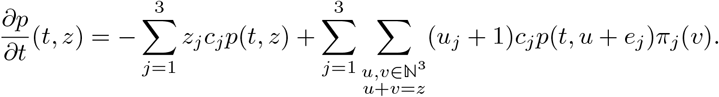

The first term is the rate at which the population departs from state *z*, and is given by the sum over all types *j* of the rate at which individuals of type *j* reproduce. The second term is the rate at which the population arrives in state *z* from another state, and can be decomposed according to the individual type whose reproduction changes the population state. Note that the descendance vector generated during the reproduction event (*v*) counts the parent when it does not die, implying that the population is formally decreased by one individual of type *j* and increased by a vector *v* during the reproduction event. In other words, if the population starts from a state *u* + *e*_*j*_ and an individual of type *j* reproduces by creating a vector *v* of descendants, the final state of the population is *u* + *e*_*j*_ − *e*_*j*_ + *v* = *u* + *v*.

Back to the derivative of *f* with respect to *t*, we use the fact that the rates *c*_*j*_ are independent of the current state of the population to re-arrange the sums and obtain:

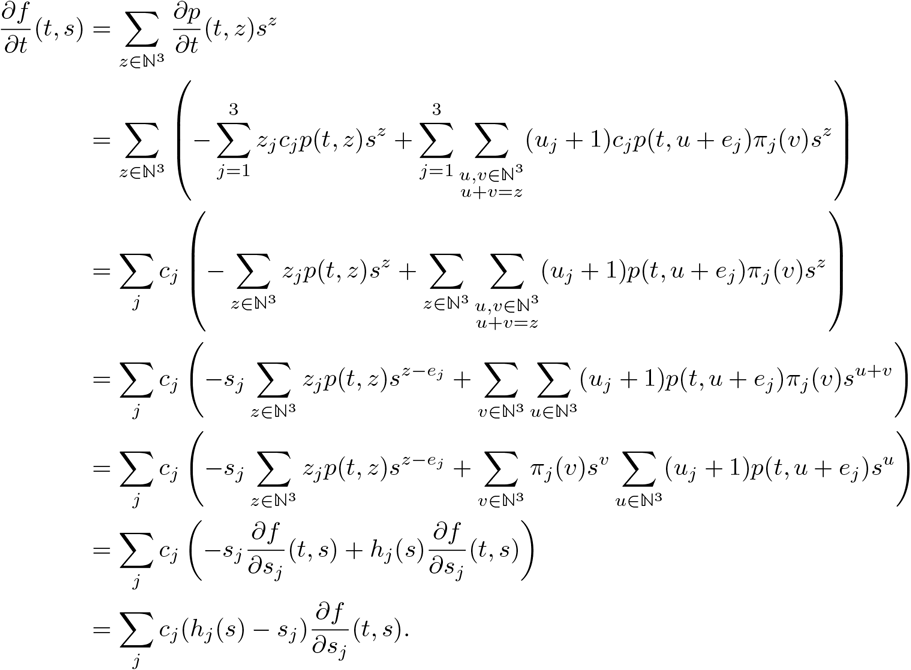

Writing *δ*_*i,j*_ = 1 if *i* = *j* and *δ*_*i,j*_ = 0 otherwise, we then obtain for the expected value:

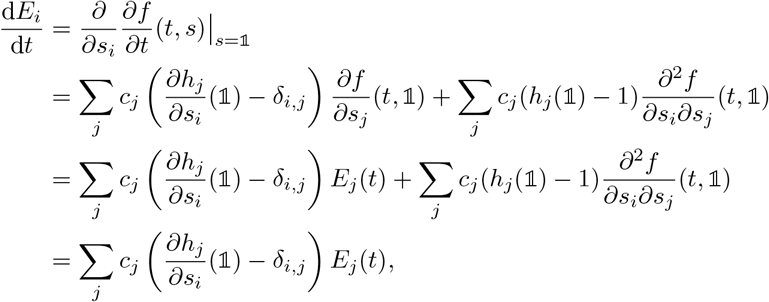

where the last equality arises from the fact that, because *h*_*j*_ is a generating function,

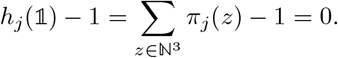

The matrix *C* we are looking for is thus defined by 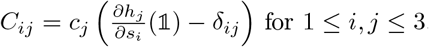.

Furthermore, we have, for all *j*,

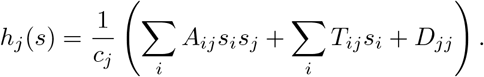

Combining the above, we arrive at

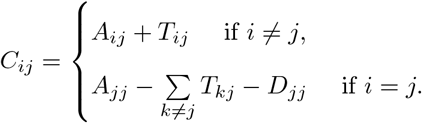

In conclusion, the matrix *C* is given by

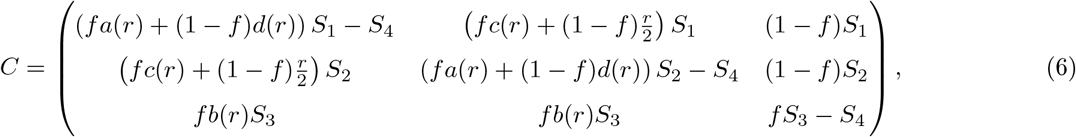

with

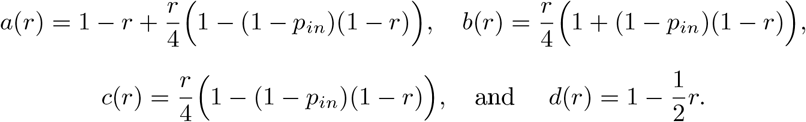

##### 7.4 Reducibility of the matrix *C* and probability of extinction of the branching process

We will use the standard notation 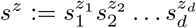 for *s* and *z* two vectors of the same dimension *d*.

Assessing the type of branching process at hand (super-, sub-, or critical) relies on the study of the eigenvalues of the matrix *C*. We use results of Sewastjanow, 1975 detailed in Pénisson, 2010 to obtain conditions on the almost-sure extinction of the process. When the matrix *C* is irreducible, the Perron-Froebenius theory of positive matrices states that it has a unique dominant eigenvalue. The branching process is then super-, sub-, or critical when this dominant eigenvalue is respectively positive, negative, or zero (Athreya and Ney, 1972, V.7.2.). In particular, the probability of extinction is equal to 1 when *ρ* ≤ 0.

In our case, the matrix *C* can be reducible (for example, when *f* = 0). In order to obtain a result on the probability of extinction in the subcritical case, we use the theory of sub-processes and of final classes. We recall below useful definitions and the principal result used (Sewastjanow, 1975).

Let (*Z*_*t*_)_*t*>0_ be a multitype branching process, with types in a finite set *K*. The equivalence r elation of *communication* is defined by: f or all states *k*_*i*_, *k*_*j*_ ∈ *K*, w e s ay that *k* _*i*_ and *k*_*j*_ *communicate*, if and only if there exist *s, t* > 0 such that

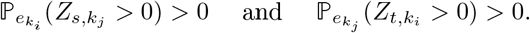

This means that there exists a time at which the probability that the population described by a branching process initiated with a single individual of type *k*_*i*_ contains an individual of type *k*_*j*_ is positive, and a time at which the probability that the population described by a branching process initiated with a single individual of type *k*_*j*_ contains an individual of type *k*_*i*_ is positive as well. If a subset 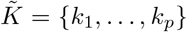 is a class for the communication equivalence relation (meaning that each state of 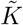 communicates with all the others but communicates with none of the states in 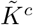), the 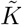 -subprocess is the process defined f or all *t* > 0 by

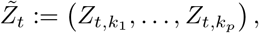

which is the vector *Z*_*t*_ from which only the coordinates of the types in the class 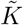 are kept. 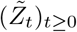 is still a branching process, and is by definition irreducible.

Let 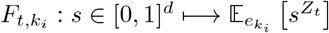 be the generating function of the process (*Z*_*t*_)_*t≥*0_ at time *t*, starting with one individual of type *k*_*i*_. 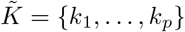 is then said to be a *final class* if it is non-empty, and satisfies the property that there exists *t* > 0 such that for all 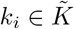 and *s* ∈ [0, 1]^*d*^, *F*_*t,k*_ (*s*) is of the form

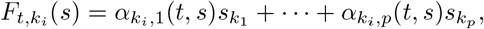

where the coefficients 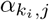 can be expressed using the coordinates *s*_*k*_ of *s* such that 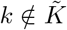. In other words, 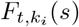 is linear in *s*_*k*_ for all 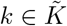. The interpretation of this property is that whenever the population starts from a single individual of type 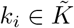, at any time *t* ≥ 0 there is one, and only one, individual of a type 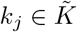 (and potentially other individuals with types in 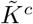). The following result gives a condition for the almost sure extinction of the process (*Z*_*t*_)_*t≥*0_ in the general case where the matrix *C* is not necessarily irreducible. Recall that the Perron’s root *ρ* of a process, when it exists, is a real eigenvalue of the matrix associated with the process such that all real parts of other eigenvalues are smaller than *ρ* (see Pénisson, 2010 Th. 1.1.7 and the following ones for a more detailed definition).

**Proposition 1** (Prop. 1.1.22 in Pénisson, 2010)

Let (*Z*_*t*_)_*t*>0_ be a continuous time Galton-Watson process, and let 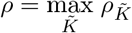 be the maximal value of the Perron’s roots of all the possible 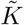-subprocesses. Then the process (*Z*_*t*_)_*t*>0_ almost surely dies out if and only if there are no final classes and *ρ⩽* 0.

Let us verify that our branching process does not contain a final class. For that, we show that the generating function of the process starting from any state has a non-zero coefficient of degree zero, and thus cannot be linear.

For any *t* > 0, *r* ∈ [0, 1]^3^ and any *j* ∈ {1, 2, 3}, we can decompose the generating function into

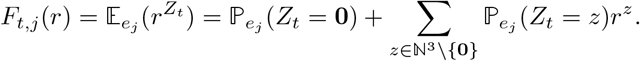

Let us prove that ℙ_*e*_*j* (*Z*_*t*_ = 0) > 0 for every *j* ∈ {1, 2, 3}. This will prove that the generating function cannot be linear for any initial type, and thus that the process does not contain any final classes.

Let *j* ∈ {1, 2, 3}, *τ*_1_ be the time of the first reproduction event, and 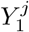 be the descendance vector created at that time. We have

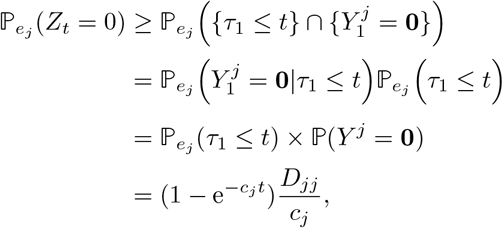

where *c*_*j*_ is the total rate of reproduction of an individual of type *j*, and *D*_*jj*_ is the rate at which an individual of type *j* reproduces and gives rise to a null vector of descendants. Hence 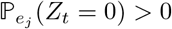 when *D*_*jj*_ > 0.

For both selection scenarii, *D*_11_ = *D*_22_ = *D*_33_ = *S*_4_. In the partial dominance selection scenario, *S*_4_ = 1, and in the overdominant selection scenario, *S*_4_ = 1 − *s*_4_. Having *D*_*jj*_ = 0 for any *j* is impossible in the first scenario and requires *s*_4_ = 1 in the second scenario, which means that the wild allele is lethal, which is not a reasonable assumption. We thus take *s*_4_ *<* 1. As a consequence, the generating function cannot be linear, and the process does not contain any final class.

The result of Proposition **??** then applies here, and the sign of the dominant eigenvalue of matrix *C* gives a condition on the almost-sure extinction of the process.

## 8 Appendices for the Results section

### 8.1 Supplementary figures

**Figure S1:**
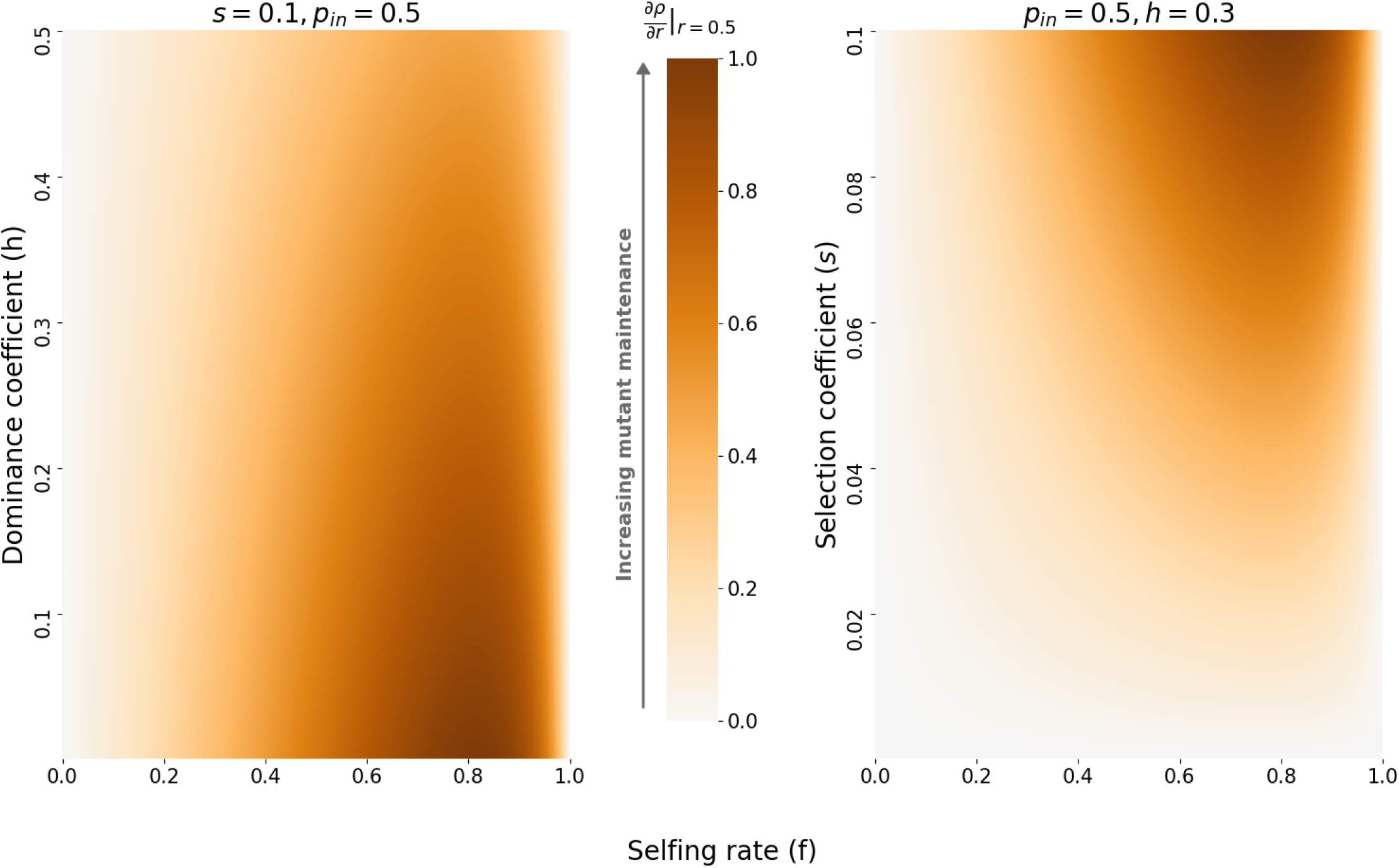
Relative variation of the derivative of the eigenvalue in the partial dominance case, for varying selfing rate *f* (x-axis), dominance coefficient *h* (y-axis, left) and selection coefficient *s* (y-axis, right). For each panel, the values of 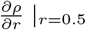 range from a minimal value, which is negative, to zero. We divided each value of the derivative by this minimum in order to plot values between 0 and 1 for every panel. This enables us to compare the impact of different parameters (*h, s* and *f*) on the sheltering effect of the mating-type locus. The darker the color, the more the mating-type locus shelters the mutation, thus promoting its maintenance.

**Figure S2:**
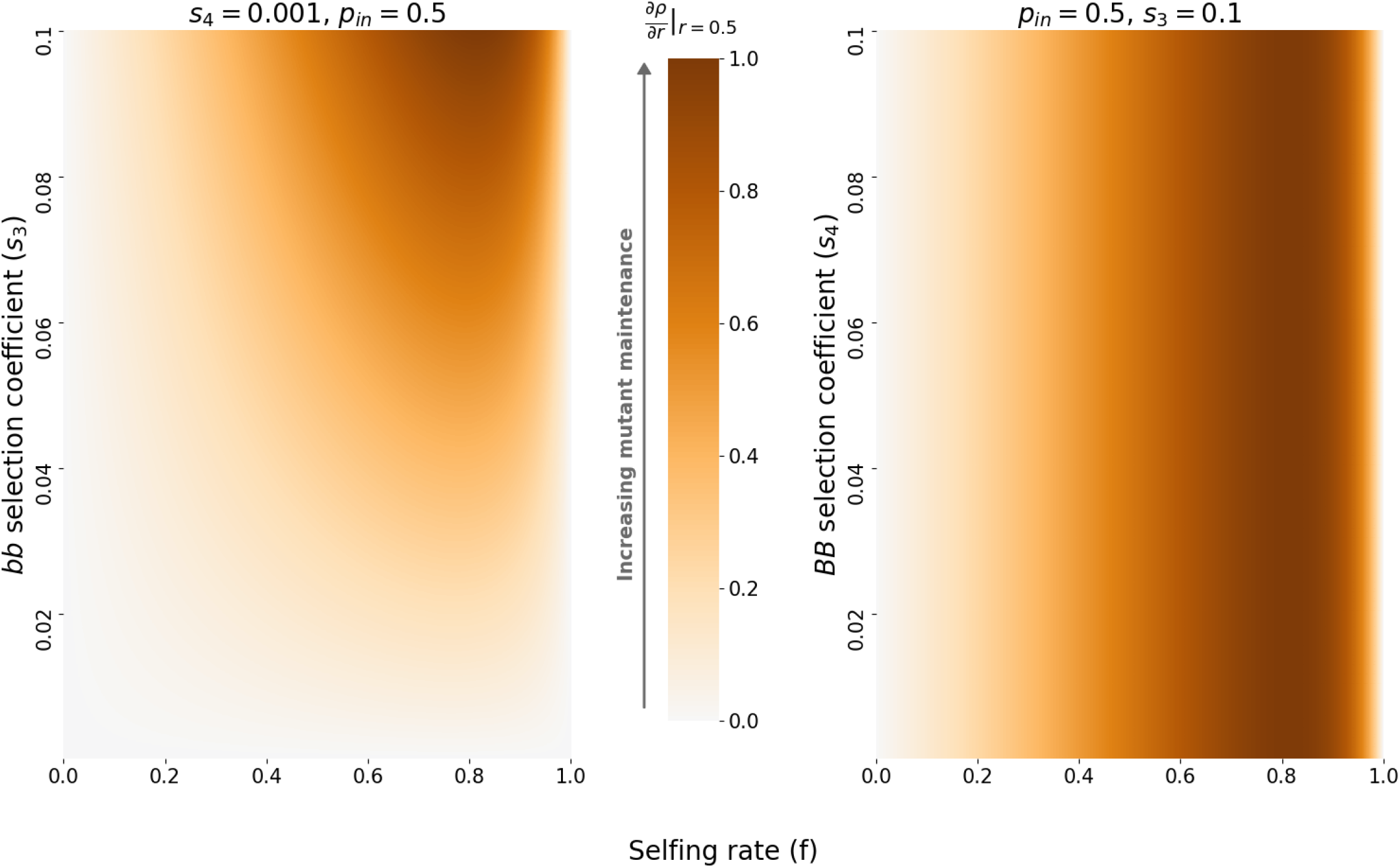
Relative variation of the derivative of the eigenvalue in the overdominance case, for varying selfing rate *f* (x-axis), and selection coefficients *s*_3_ (y-axis, left) and *s*_4_ (y-axis, right), with *s*_3_ > *s*_4_. For each panel, the values 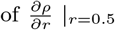 range from a minimal value, which is negative, to zero. We divided each value of the derivative by this minimum in order to plot values between 0 and 1 for every panel. This enables us to compare the impact of different parameters (*s*_3_, *s*_4_ and *f*) on the sheltering effect of the mating-type locus. The darker the color, the more the mating-type locus shelters the mutation, thus promoting its maintenance.

**Figure S3:**
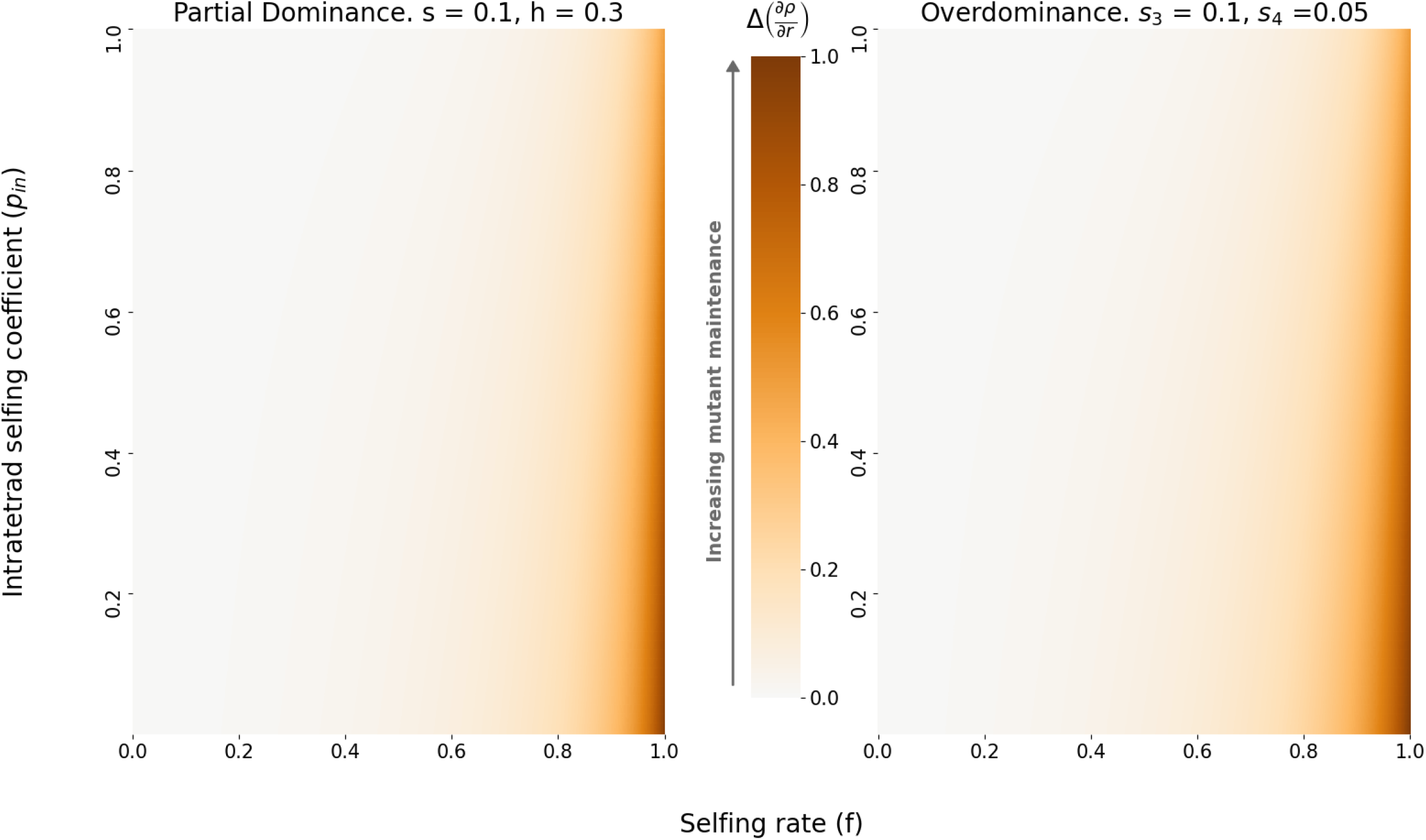
Difference between the dominant eigenvalue derivative at *r* = 0.5 and at 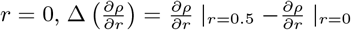. The left panel shows the partial dominance case, the right panel shows the overdominance case, for varying selfing rate *f* (x-axis), and intra-tetrad selfing rate (y-axis). The difference is always positive, with both derivative being negative (see App. **??** and App. **??**). This means that the absolute value of the derivative at *r* = 0 is always greater than the absolute value of the derivative at *r* = 0.5. The darker the color, the larger the difference between the two derivatives.

**Figure S4:**
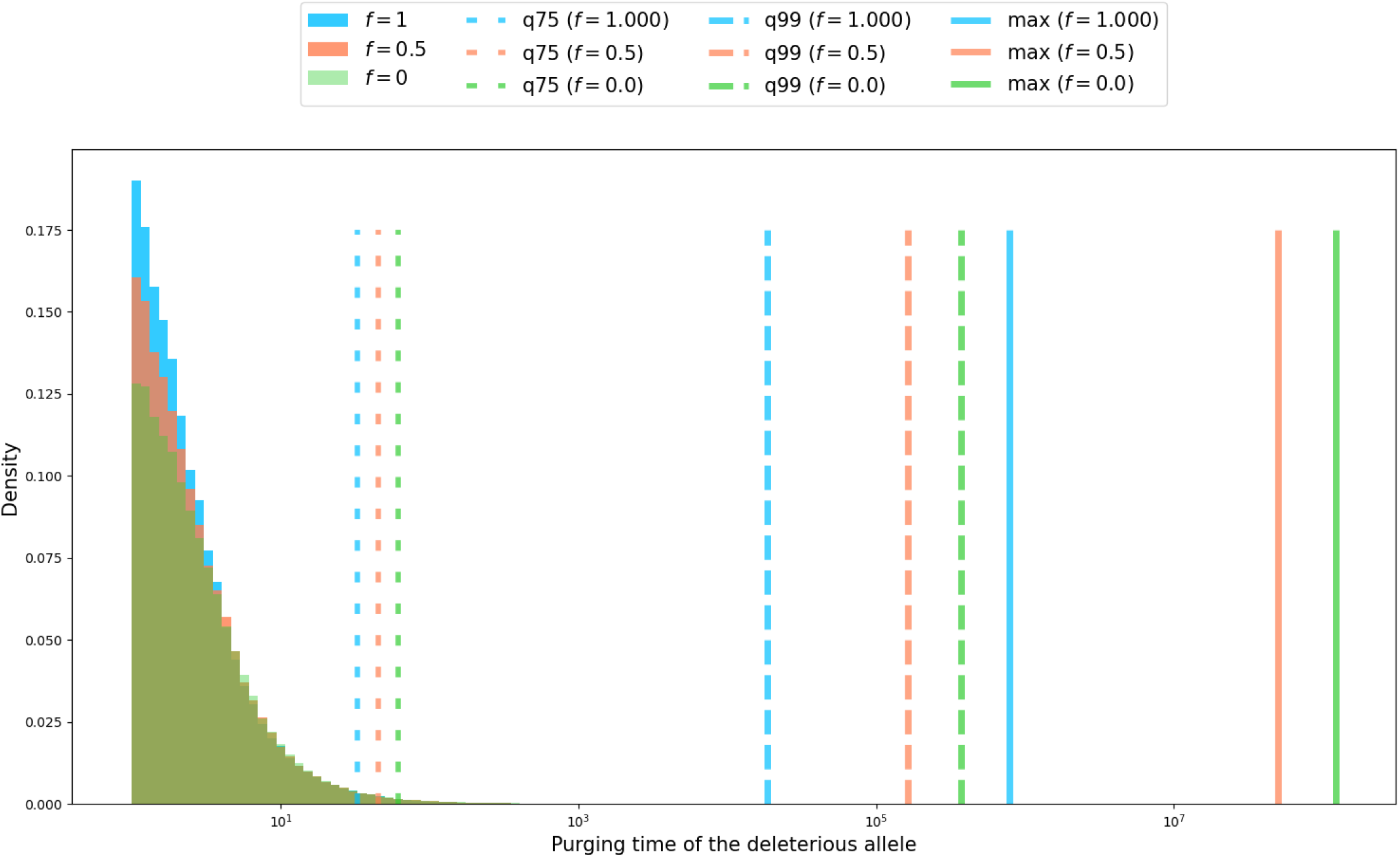
Empirical distribution of the deleterious allele purging time for the partial dominance scenario. A total of 100,000 simulations were run, with *s* = 0.1, *h* = 0.1, *r* = 0.1, *p*_*in*_ = 0.5, starting from one heterozygous individual (*X*_0_ = (1, 0, 0)), and for three values of the selfing rate (*f* = 0 in green, *f* = 0.5 in red and *f* = 1 in blue). The respective values for *ρ* are *ρ* = − 0.0100, *ρ* = − 0.0157 and *ρ* = − 0.0818. The x-axis is log-scaled. The large-dotted lines represent the 75^*th*^ percentile (*q75*), the dashed lines indicate the 99^*th*^ percentile (*q99*), and solid lines the maximum value (*max*) of the purging time. Maximum values are several order of magnitudes higher than the 75th percentile of the empirical distribution of the purging time.

**Figure S5:**
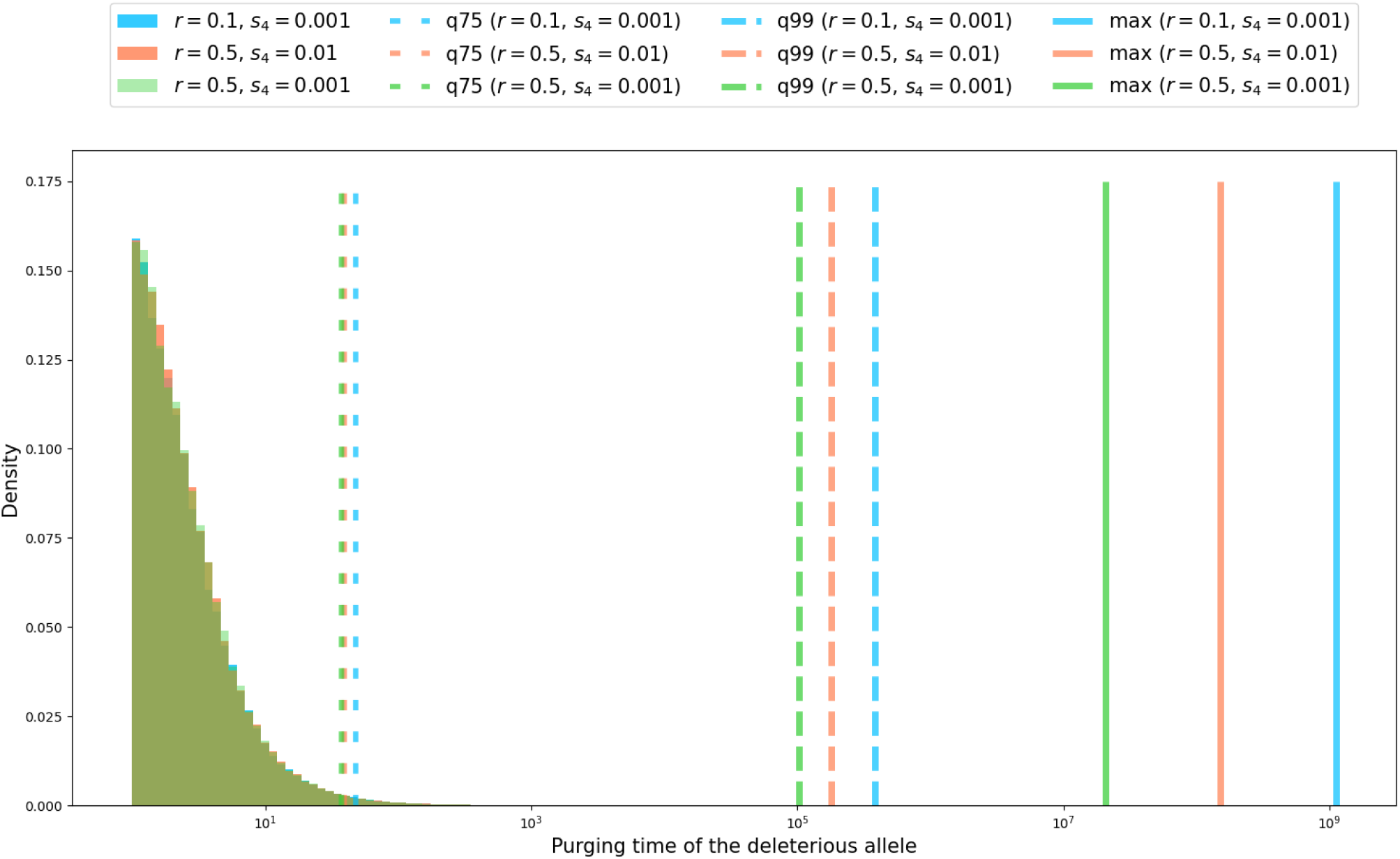
Empirical distribution of the deleterious allele purging time for the overdominance scenario. A total of 100,000 simulations were run, with *s*_3_ = 0.1, *f* = 0.5, *p*_*in*_ = 0.5, starting from one heterozygous individual (*X*_0_ = (1, 0, 0)), for several values of the recombination rate and of the selection coefficient *s*_4_ (*r* = 0.1, *s*_4_ = 0.001 in blue, *r* = 0.5, *s*_4_ = 0.01 in red, and *r* = 0.5, *s*_4_ = 0.001 in green). The respective values for *ρ* are *ρ* = − 0.0052, *ρ* = −0.0129 and *ρ* = − 0.0219. The parameters were chosen so that the process is sub-critical and thus the purging time is almost surely finite. The x-axis is log-scaled. The large-dotted lines represent the 75^*th*^ percentile (*q75*), the dashed lines indicate the 99^*th*^ percentile (*q99*), and solid lines the maximum value (*max*) of the purging time. Maximum values are several order of magnitudes higher than the 75th percentile of the empirical distribution of the purging time.

**Figure S6:**
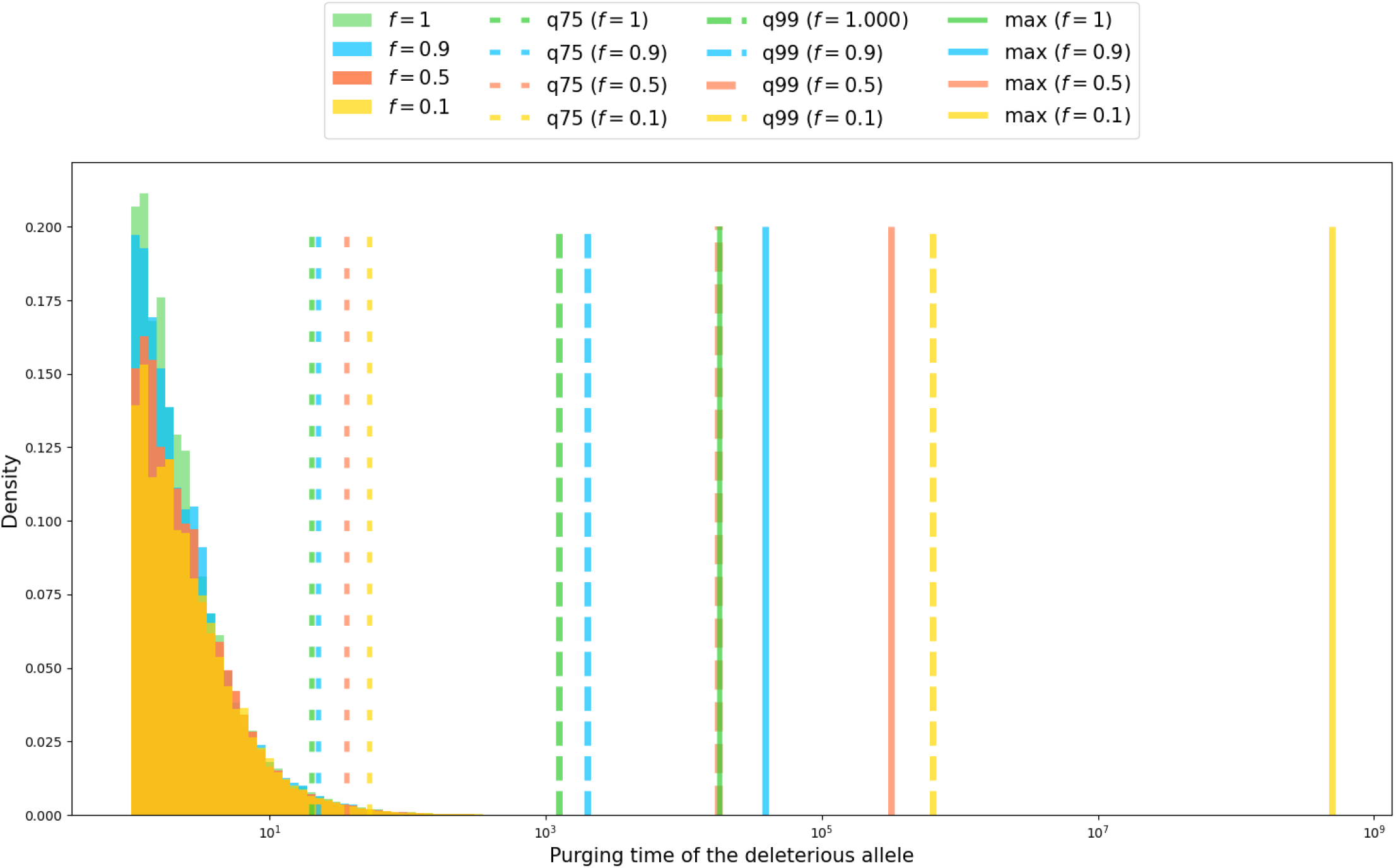
Empirical distribution of the deleterious allele purging time for the overdominant scenario. A total of 100,000 simulations were run, with *s*_3_ = 0.5, *p*_*in*_ = 0.5, *s*_4_ = 0.01, *r* = 0.4, starting from one heterozygous individual (*X*_0_ = (1, 0, 0)), for four values of the selfing rate (*f* = 0.1 in yellow, *f* = 0.5 in red, *f* = 0.9 in blue, *f* = 1 in green). The respective values for *ρ* are *ρ* = − 0.0035, *ρ* = − 0.0713, *ρ* = − 0.1905 and *ρ* = − 0.25. Parameters were chosen so that the process is sub-critical and thus the purging time is almost surely finite. The x-axis is log-scaled. The large-dotted lines represent the 75^*th*^ percentile (*q75*), dashed lines indicate the 99^*th*^ percentile (*q99*), and solid lines the maximum value (*max*) of the purging time. Maximum values are several order of magnitudes higher than the 75th percentile of the empirical distribution of the purging time. Note that the selection coefficient for *bb* homozygotes is high (*s*_3_ = 0.5).

### 8.2 The dominant eigenvalue, its sign and its derivative: partial dominance scenario

#### 8.2.1 Determination of the dominant eigenvalue

The eigenvalues computed with Mathematica (Wolfram Research, 2015) are, for the partial dominance case,

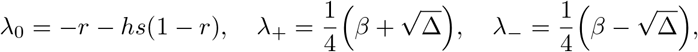

where

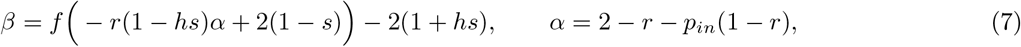

and

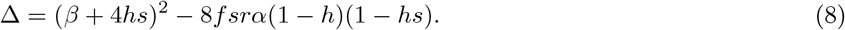

It is straightforward to see that *λ*_+_ > *λ*_−_.

Let us prove that we also have *λ*_+_ > *λ*_0_. We used Geogebra to assist us in the calculations.

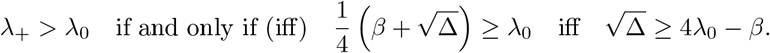

If 4*λ*_0_ − *β* ≤ 0, the last inequality if straightforward, as 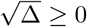.

Let us study the sign of 4*λ*_0_ − *β*. We define *P* (*r*) := 4*λ*_0_ − *β* = *a*_2_*r*^2^ + *a*_1_*r* + *a*_0_, with

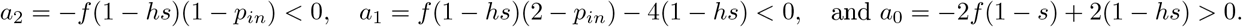

*P* is a second-order polynomial, with negative quadratic coefficient and positive coefficient of order zero (because 1 − *hs* > 1 − *s* > *f* (1 − *s*)). Thus, *P* admits two roots, one negative and one positive. We denote the positive root by *r*_*P*_. For *r* ∈ [0, *r*_*P*_], we have *P* (*r*) ≥ 0, and for *r* > *r*_*P*_, we have *P* (*r*) *<* 0. Consequently, we readily obtain that when *r* > *r*_*P*_, *λ*_+_ > *λ*_0_.

Let us now consider the case *r* ∈ [0, *r*_*P*_]. For such an *r*, using that 4*λ*_0_ − *β* ≥ 0, we can write that

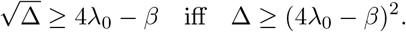

Let us write *Q*(*r*) := (4*λ*_0_ − *β*)^2^ − Δ = *b*_2_*r*^2^ + *b*_1_*r* + *b*_0_, with

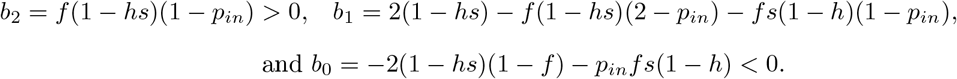

*Q* is a second-order polynomial, with positive quadratic coefficient, and negative coefficient of order 0. Hence, *Q* admits two roots, one negative, and one positive. We denote the positive root by *r*_*Q*_. In order to prove that *λ*_+_ > *λ*_0_, we have to prove that *Q*(*r*) ≤ 0 when *P* (*r*) > 0, *i.e*. when *r* ∈ [0, *r*_*P*_]. As *Q*(0) *<* 0 and *Q* has only one positive root, proving that *Q*(*r*_*P*_) *<* 0 will imply that *Q*(*r*) ≤ 0 for *r* ∈ [0, *r*_*P*_]. Let us prove that *Q*(*r*_*P*_) *<* 0. Noting that the quadratic coefficients of *P* and *Q* are the opposites of one another, we use the equation *P* (*r*_*P*_) = 0 to obtain

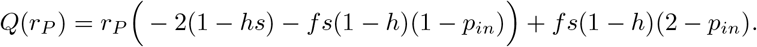

Seeing *Q*(*r*_*P*_) as an affine function of *r*_*P*_, we obtain that the function *r*_*P*_ ⟼ *Q*(*r*_*P*_) admits a unique root, which is positive, and that we will denote by 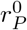:

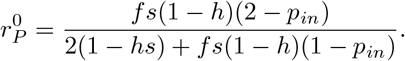

We wish to prove that 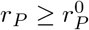, as it implies that *Q*(*r*_*P*_) ≤ 0. Having 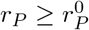 is equivalent to having 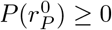, as *r*_*P*_ is the unique positive root of *P* and *P* (0) ≥ 0. Consequently, it only remains to prove that 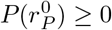, which is equivalent to

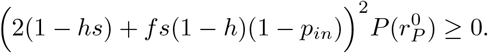

To obtain this result, an efficient way is to consider the left-hand term as a polynomial in *p*_*in*_. Let us write 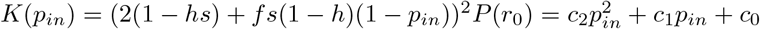 with

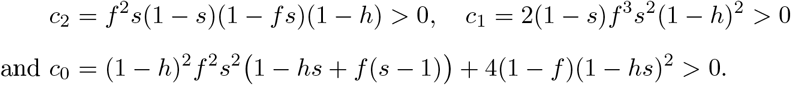

*K* is thus a second-degree polynomial in *p*_*in*_, with a positive quadratic coefficient, and a minimum reached for a negative value (minimum reached at −*c*_1_/(2*c*_2_) *<* 0). *K* is thus monotonic for positive abscissa, and the coefficient of order zero is positive. Consequently, for all *p*_*in*_ ≥ 0, we have *K*(*p*_*in*_) ≥ 0. We have then *P* (*r*_0_) ≥ 0, which concludes the proof that *λ*_+_ ≥ *λ*_0_.

Based on the result we just obtained, from now on we write *ρ* = *λ*_+_.

#### 8.2.2 Sign of the dominant eigenvalue

We prove that *ρ <* 0, except when *s* = 0, or when *h* = 0 and *r* = 0, in which cases *ρ* = 0. Recall the notation *α, β* from (**??**) and Δ from (**??**).

First, considering that *r* ∈ [0, 1] and *p*_*in*_ ∈ [0, 1], we have 0 *< α <* 2, which leads to *β <* 0.

When *s* = 0 or (*r, h*) = (0, 0), Δ = *β*^2^, which gives, as 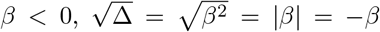. We then have 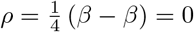.

Let us now consider the case where *s* ≠ 0 and (*r, h*) ≠ (0, 0). We have

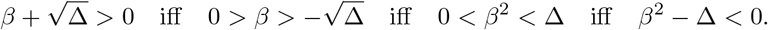

Moreover, *β*^2^ − Δ = −16*f* (1 − *s*)*hs* + 8*frα*(1 − *hs*)*s* + 16*hs*. The sign of *ρ* is thus the sign of *fh*(2 − *rα*)*s* + 2*h*(1 − *f*) + *rαf*, which is an affine function of *s*. The slope and intercept of this function are both non-positive when (*r, h*) ≠ 0 or *s* ≠ 0, which gives *ρ <* 0 in those cases.

#### 8.2.3 Derivative of the dominant eigenvalue

The derivative of *ρ* is

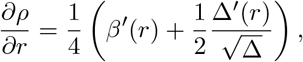

Evaluating this derivative at *r* = 0.5, and using that *β*′(0.5) = −*f* (1 − *hs*), we obtain

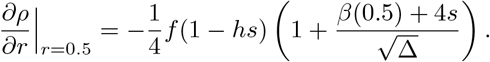

Simple calculations lead to 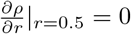 when *f* = 0, or *s* = 0, or *s* = 1, or *h* = 1, or *f* = 1 and 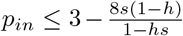.

In the latter case, whether the inequality is verified or not determines the sign of Δ and therefore the value of 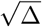, which is either equal to *β*(0.5) + 4*s* or −(*β*(0.5) + 4*s*). The derivative is then either equal to zero or strictly negative.

For the rest of this paragraph, we study the sign of the derivative when none of the above cases is met.

Let us write *γ* = *β*(0.5) + 4*s*. If *γ* ≥ 0, we readily obtain 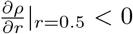. Let us then assume that *γ <* 0. In this case, we have

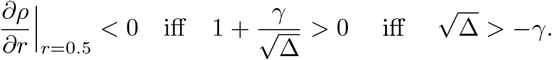

As −*γ* > 0, this comes down to

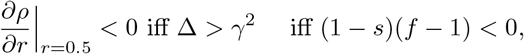

which is indeed satisfied.

In conclusion, we have shown that, in the general case,

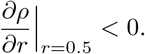

We also compute the derivative at *r* = 0. We have *β*(0) = 2*f* (1 − *s*) − 2(1 + *hs*), *β*′(0) = −*f* (1 − *hs*)(2 − *p*_*in*_), Δ(0) = (2*f* (1 − *s*) − 2(1 − *hs*))^2^, and Δ′(0) = 2*β*′(0)(*β*(0) + 4*hs*) − 8*fs*(1 − *h*)(1 − *hs*)(2 − *p*_*in*_).

After simplification, this gives

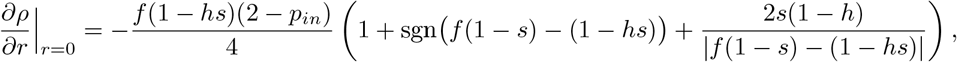

with sgn (*f* (1 − *s*) − (1 − *hs*)) is equal to 1 (respectively to −1) when *f* (1 − *s*) − (1 − *hs*) is positive (resp. negative).

We obtain immediately that

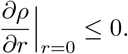

### 8.3 The dominant eigenvalue, its sign and its derivative: overdominance scenario

#### 8.3.1 Determination of the dominant eigenvalue

The eigenvalues computed with Mathematica (Wolfram Research, 2015) for the overdominant case are

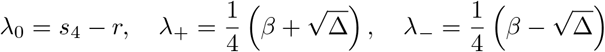

with

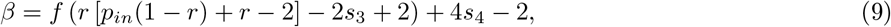

and

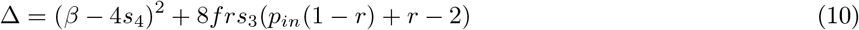

Here again, we obviously have *λ*_+_ > *λ*_−_.

We follow the same method as in the partial dominance case to prove that *λ*_+_ > *λ*_0_.

We have

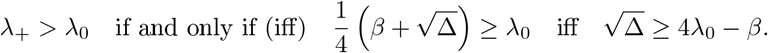

If 4*λ*_0_ − *β* ≤ 0, the last inequality if straightforward, as 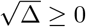. Let us thus study the sign of 4*λ*_0_ − *β*. Let us define the function *P* by *P*(*r*) := 4*λ*_0_ − *β* = *a*_2_*r*^2^ + *a*_1_*r* + *a*_0_, with

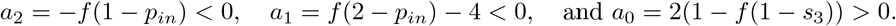

*P* is a second-order polynomial, with a negative quadratic coefficient and a positive coefficient of order 0. Hence *P* admits two roots, one which is negative and one which is positive. We denote the positive root by *r*_*P*_. For *r* ∈ [0, *r*_*P*_], we have *P* (*r*) ≥ 0, and for *r* > *r*_*P*_, we have *P* (*r*) *<* 0. Consequently, we readily obtain that when *r* > *r*_*P*_, the conclusion follows.

Let us now consider *r* ∈ [0, *r*_*P*_]. For such an *r*, as 4*λ*_0_ − *β* ≥ 0, again we have

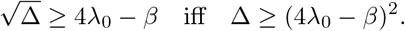

Let us define the function *Q* by *Q*(*r*) := (4*λ*_0_ − *β*)^2^ − Δ = *b*_2_*r*^2^ + *b*_1_*r* + *b*_0_, with

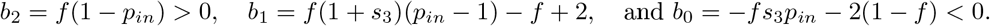

*Q* is a second-order polynomial, with positive quadratic coefficient, and negative coefficient of order 0. Hence, *Q* admits two roots, one negative and one positive. We denote the positive root by *r*_*Q*_. In order to prove that *λ*_+_ > *λ*_0_, we have to prove that *Q*(*r*) ≤ 0 when *P* (*r*) > 0, *i.e*. when *r* ∈ [0, *r*_*P*_]. As *Q*(0) *<* 0 and *Q* has only one positive root, proving that *Q*(*r*_*P*_) *<* 0 will imply that *Q*(*r*) ≤ 0 for *r* ∈ [0, *r*_*P*_]. Let us prove that *Q*(*r*_*P*_) *<* 0. Noting that the quadratic coefficients of *P* and *Q* are the opposites of one another, we use the equation *P* (*r*_*P*_) = 0 to obtain

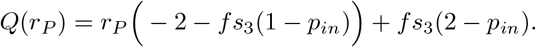

Seeing *Q*(*r*_*P*_) as an affine function of *r*_*P*_, we obtain that the function *r*_*P*_ ⟼ *Q*(*r*_*P*_) admits a unique root, which is positive, and that we will denote by 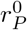:

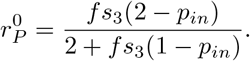

We wish to prove that 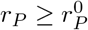, as it implies that *Q*(*r*_*P*_) ≤ 0. Having 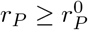 is equivalent to having *P* (*r*_0_) ≥ 0, as *r*_*P*_ is the unique positive root of *P* and *P* (0) ≥ 0. There is thus left to prove that 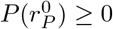, which is equivalent to

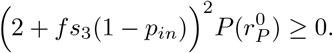

To obtain this result, an efficient way is to consider the left-hand term as a polynomial in *p*_*in*_.

Let us write *K*(*p*_*in*_) = (2 + *fs*_3_(1 − *p*_*in*_))^2^*P* (*r*^0^) = *c*_2_*p*^2^ + *c*_1_*p*_*in*_ + *c*_0_, with

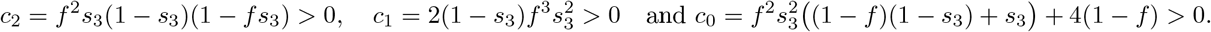

*K* is thus a second-degree polynomial in *p*_*in*_, with a positive quadratic coefficient and positive coefficient of order 0, that reaches its minimum for a negative value (minimum reached at −*c*_1_/(2*c*_2_) *<* 0). Consequently, for all *p*_*in*_ ≥ 0, we have *K*(_*Pin*_) ≥ 0. We have then 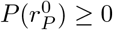, which concludes the proof that *λ*_+_ ≤ *λ*_0_.

Based on the result we just obtained, from now on we write *ρ* = *λ*_+_.

#### 8.3.2 Sign of the dominant eigenvalue

In this selection scenario, *ρ* is not of constant sign.

The condition for *ρ* ≥ 0 is

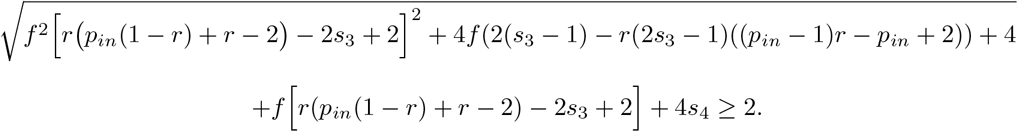

We compute the dominant eigenvalue and study its sign for simple cases, and then use a numerical approach to complete the analysis (Figure **??**). Under complete intra-tetrad selfing (*f* = 1, *p*_*in*_ = 1), we have *ρ* = *s*_4_ − *r*/2 if *s*_3_ ≥ *r*/2 and *ρ* = *s*_4_ − *s*_3_ if *s*_3_ *< r*/2. As *s*_4_ ≤ *s*_3_, the condition to have *ρ* ≥ 0 reduces to *r* ≤ 2*s*_4_. This is consistent with the results of Antonovics and Abrams, 2004, as the authors set *s*_3_ = 1 and thus obtain *ρ* = *s*_4_ − *r*/2. Under complete selfing (*f* = 1), if *r*(2 − *r* − *p*_*in*_(1 − *r*)) − 2*s*_3_ ≥ 0, then *ρ* = *s*_4_ − *s*_3_ ≤ 0. This shows that the value of the dominant eigenvalue, and thus the dynamics of the process, depends only on the selection strength when the recombination rate *r* exceeds a certain threshold. Moreover, this threshold depends only on the selection coefficient for homozygous deleterious (*s*_3_), and on the probability of intra-tetrad mating (*p*_*in*_). This threshold appears on the bottom panels of Figure **??**. Under complete outcrossing (*f* = 0), we have *ρ* = *s*_4_ ≥ 0. When the mutation is completely linked to a mating-type allele (*r* = 0), we have *ρ* = *s*_4_ ≥ 0. When the mutation is neutral (*s*_3_ = 0, implying *s*_4_ = 0 as well), we have *ρ* = 0. Finally, when *BB* homozygotes are not disfavored (*s*_4_ = 0), we have *ρ <* 0. Indeed, in this case, *β* = *fr*[*p*_*in*_(1 − *r*) + *r* − 2] − 2*fs*_3_ + 2(*f* − 1) *<* 0. We thus have

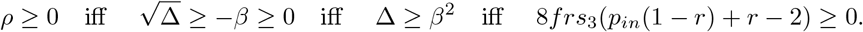

But we trivially have *p*_*in*_(1 − *r*) + *r* − 2 ≤ 0, and so the condition is not met and *ρ <* 0.

#### 8.3.3 Derivative of the dominant eigenvalue

The derivative of the largest eigenvalue *ρ* is

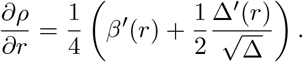

Moreover, we have

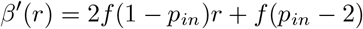

and

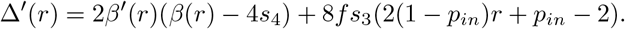

Evaluating these quantities at *r* = 0.5, we obtain *β*′ (0.5) = −*f*, and so

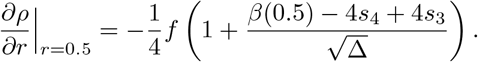

Simple calculations lead to 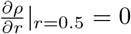 when *f* = 0, or *s*_3_ = 0, or *f* = 1 and *p*_*in*_ ≤ 3 − 8*s*_3_.

For the rest of this paragraph, we study the sign of the derivative when none of the above cases is met.

Let us write *γ* = *β*(*r* = 0.5) − 4*s*_4_ + 4*s*_3_. If *γ* ≥ 0, we readily obtain that 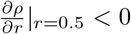. Let us then assume that *γ <* 0. In this case, we have

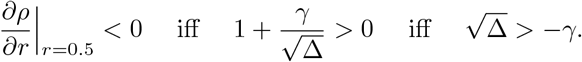

As −*γ* > 0,

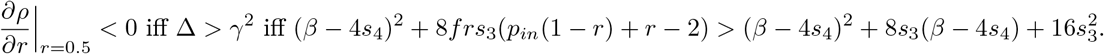

After some simplifications, we obtain

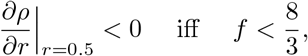

which is always satisfied as *f* ∈ [0, 1].

In conclusion, we have shown that

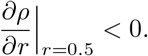

We also compute the derivative at *r* = 0. We have *β*(0) = 2*f* (1 − *s*_3_) + 4*s*_4_ − 2, *β*′ (0) = −*f* (2 − *p*_*in*_), Δ(0) = (2*f* (1 − *s*_3_) − 2)^2^, and Δ′ (0) = 2*β*′ (0)(*β*(0) − 4*s*_4_) − 8*fs*_3_ (2 − *p*_*in*_).

After simplification, this gives

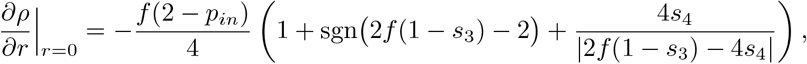

with sgn (2*f* (1 − *s*_3_) − 2) is equal to 1 (respectively to −1) when 2*f* (1 − *s*_3_) − 2 is positive (resp. negative). We obtain immediately that

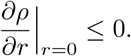

